# Motor learning- and consolidation-related resting state fast and slow brain dynamics across wake and sleep

**DOI:** 10.1101/2023.07.27.550839

**Authors:** Liliia Roshchupkina, Vincent Wens, Nicolas Coquelet, Charline Urbain, Xavier de Tiege, Philippe Peigneux

## Abstract

Learning and consolidation of motor skills dynamically evolve both online during practice and offline after training. We investigated, using magnetoencephalography, the neural dynamics underpinning motor learning and its consolidation in relation to sleep during resting-state periods shortly after the end of learning (short-term boost window, within 30 min) and at more delayed time scales (silent 4h and next day 24h windows) with an intermediate nap or wakefulness after the boost window. Resting-state neural dynamics in brain networks were investigated at fast (sub-second) and slower (supra-second) timescales using Hidden Markov modelling (HMM) and resting-state functional connectivity (rsFC), respectively, as well as their relationship with the evolution of motor performance. HMM results show that fast dynamic activities in a Temporal/Sensorimotor state network predict individual motor performance achievements, suggesting a trait-like association between rapidly recurrent neural patterns and motor behaviour. Short, post-training re-exposure to the task modulated fast and slow network characteristics during the boost but not in the silent window. These short practice-related induction effects were observed again on the next day, to a reduced extent as compared to the boost window. Daytime naps did not significantly modulate memory consolidation both at behavioural and neural levels. These results emphasise the critical role of the transient boost window in motor learning and subsequent memory consolidation processes and provide further insights into the relationship between the multiscale neural dynamics of brain networks, motor learning, and consolidation.

## INTRODUCTION

Motor learning (ML) is an essential, complex process allowing one to acquire, maintain and adapt motor-based behavioural responses to ever-changing environmental demands, eventually enabling efficient and adaptive functioning. To learn a new motor skill (e.g., knitting or crocheting, writing, dancing …), one must repeatedly practice movements to achieve swift and accurate performance. ML not only rapidly develops online during task practice but also continues to unfold at a slower but significant rate, leading to additional performance gains after the end of the actual learning episode. This process, known as the offline post-training consolidation phase, unfolds non-linearly across critical time windows after training [1], [2]. As compared to the end of the ML episode, there is a significant performance gain at retest 5 to 30 minutes after the end of practice [3], [4]. However, performance enhancement within this so-called performance boost window period is only transient, as it remains at end-of-learning levels when tested a few hours later (i.e., a silent window), then improves again on the following day [3], [4]. Performance gains achieved at the early, short-lived boost phase were found predictive of offline performance improvement 48h later [4], suggesting the functional relevance of immediate post-training periods for the rapid coordination and reorganisation of neural networks supporting ML and its consolidation on the long term [1]–[4]. Additionally, post-learning sleep was proposed to contribute to the consolidation of motor memories. Amongst others, supporting evidence comes from studies showing enhanced motor performance after a post-learning nap episode [5]–[8] and robust motor performance benefits after overnight sleep [9] (but see, e.g., [10] for discrepant results). Noticeably, studies on the dynamics of post-learning nap long-term consolidation were found inconsistent, with post-nap motor performance gains either found to vanish [11] or persist [5] after a full night of sleep. Available data suggest that sleep-dependent motor memory consolidation effects are contingent upon multiple boundary conditions, including the specificities of the practiced motor task, the contributing neural substrates and the intervening sleep period components [12], which may explain discrepancies in the outcome of sleep ML studies.

The neuroanatomical underpinnings of ML and their functional interactions at the brain network level are well documented [13]–[16]. Functional connectivity measures allow investigating the functional brain network architecture both during actual ML practice (i.e., task-based connectivity) and during off-task, non-directed resting-state periods before and after the learning episode (i.e., resting-state (RS) connectivity). Previous studies demonstrated online and offline changes in ML-related neural dynamics (NDs) on a timescale ranging from seconds to minutes [17]–[20]. Online neural network features during ML (e.g., flexibility [21], local path length, connectivity strength and nodal efficiency [22]) reconfigure over practice and can be predictors of future learning developments [21]. Similarly, functional network connectivity derived from offline RS measurements prior to a motor task was shown to predict individual ML abilities [20], [23], and post-learning RS features were found to reflect task-induced plasticity [17], [22]. An asset of offline RS neuroimaging is to allow investigating learning-related functional changes in the brain architecture uncontaminated, e.g., by actual task-performance bias or task-induced movement artefacts.

Resting-state networks (RSNs) not only feature rich spatiotemporal dynamics [24]–[27] but are also shown to exhibit frequency-specific characteristics [28]–[30]. Consequently, RS neuroimaging can offer valuable insights into the neural plasticity mechanisms underlying ML and its consolidation. It also allows investigating the different timescales at which brain networks are being coordinated and reorganised in response to changing cognitive demands. While RSNs connectivity evolves over slow, supra-second timescale dynamics [31]–[33], Baker and colleagues [34] showed, using hidden Markov modelling (HMM) of band-limited power envelopes in magnetoencephalography (MEG) recordings, that activity patterns within RSNs change much more rapidly than previously thought. Indeed, they and later studies [35]–[40] reported discrete transient (100–200 msec duration) brain states repeatedly reoccurring over time in RS-related neural activity, corresponding to activation/deactivation power patterns in well-known RS networks [33], [41], [42]. HMM outcomes may provide support to the hypothesis that neurocognitive networks adapt to the rapidly changing computational demands of cognitive processing [43] through rapid reorganisation and coordination mechanisms operating at the sub-second timescale [44]. Furthermore, exploring the synchronisation of brain networks within specific frequency bands at supra-seconds scale may offer deeper insights into ML-related mechanisms. Therefore, taking advantage of the excellent spatiotemporal resolution of MEG, combining HMM with RSN connectivity analysis opens prospects to investigate the neural plasticity dynamics underlying ML and its consolidation at various temporal scales.

In the present MEG study, we investigated the spontaneous, multiscale NDs underlying ML and its consolidation at critical short-term and delayed RS periods, modulated by the availability of post-training diurnal sleep and task re-exposure. We specifically analysed the fast (sub-second) activation dynamics of network states and the slower (supra-second) modulations that give rise to functional connectivity, investigating their interrelationships and associations with motor performance over time. We hypothesised that ML would modulate RS NDs at both fast and slow timescales, reflecting neural bases of learning/consolidation process. Further, we posited that a brief re-exposure to the learned motor task (induction effect) would reinstate and reinforce activity in ML-related networks, and that delayed effects on RS NDs will be modulated by an intermediate sleep period during a daytime nap.

## METHODS

### Participants

Thirty-four young, healthy right-handed participants gave written informed consent to participate in this study approved by the Ethics Committee of the CUB-ULB Erasme Hospital (Brussels, Belgium) (Ref: P2016/553; CCB: B406201630539). Musicians and professional typists were excluded to avoid bias due to high baseline motor skill proficiency. Four participants were excluded based on preliminary analyses (3 due to insufficient motor performance; 1 due to a corrupted MEG signal). Data for 30 participants (10 females; mean age = 23.1 ± 2.7 yrs, range 18 - 29) are reported. They received monetary compensation for their participation.

### Procedure

To avoid hormonal influence on motor learning abilities, all female participants were tested during the second week of their menstrual cycle [45]. Caffeine-containing drinks, food, soda, and any other stimulants were prohibited for 12 h before testing.

The experimental setting and task are illustrated in Figure 1. Participants arrived at the laboratory at 8 am to be prepared for MEG recordings. The first (baseline) 5-minute resting-state (RS 1) recording session took place in the MEG-shielded room, seated with eyes open and focused on a fixation cross on the screen. Then, they filled in visual analogue scales (VAS) for fatigue [46] and sleepiness to control for potential drowsiness and fatigue effects. Immediately after, they were trained on a 5-element Finger Tapping Task (FTT [4] adapted from [47]). In the FTT (Figure 1), each finger corresponds to one digit (from 1 = little finger to 4 = index), and participants are instructed to continuously reproduce with their non-dominant hand a fixed 5-element sequence of finger movements (4-1-3-2-4; permanently displayed on the computer screen) as fast and accurately as possible for 30 seconds (i.e., 1 block). They were familiarised with the task during 2 FTT demo blocks before performing the experimental learning session (LS) that comprised twenty 30-second FTT blocks (separated by 20-second breaks). Twenty minutes after the end of FTT practice (boost window session), they filled in again VAS for fatigue and sleepiness then were administered a first 5-min RS session (RS 2), followed by performance testing on 2 FTT blocks and then, again, a RS session (RS 3). The rationale for having RS recordings before and after behavioural testing was to investigate spontaneous RS activity after a consolidation period and then after the reactivation of the motor network due to the short FTT practice at testing, i.e., induced by immediately preceding motor practice. Participants could then enjoy free time and lunch. Afterwards, participants were told that they were randomly assigned either to a Wake (n = 16, five females) or Nap (n = 18, six females) condition. Participants from the Wake group stayed awake for the next 1.5 h, while the Nap group could benefit from a 90-min nap. Four hours after the end of the learning session (silent window session), all participants underwent a second testing block identical to the first one, with RS before (RS 4) and immediately after (RS 5) a short behavioural testing (2 FTT blocks). Participants were then kept in the laboratory and allowed to rest, communicate with the researchers, or watch movies until the evening. Around 9 pm, they were prepared for overnight High-density EEG (HD-EEG) sleep recordings. Bedtime was set to 11 pm ± 30 min, and participants did sleep for about 7h. After waking up on the next day, they were prepared for the MEG recordings and administered a final session (next day window) with RS before (RS 6) and after (RS 7) a short behavioural testing (2 FTT blocks). Finally, participants underwent a structural MRI scan for the sake of MEG source reconstruction (see MEG methods).

**Figure 1.**
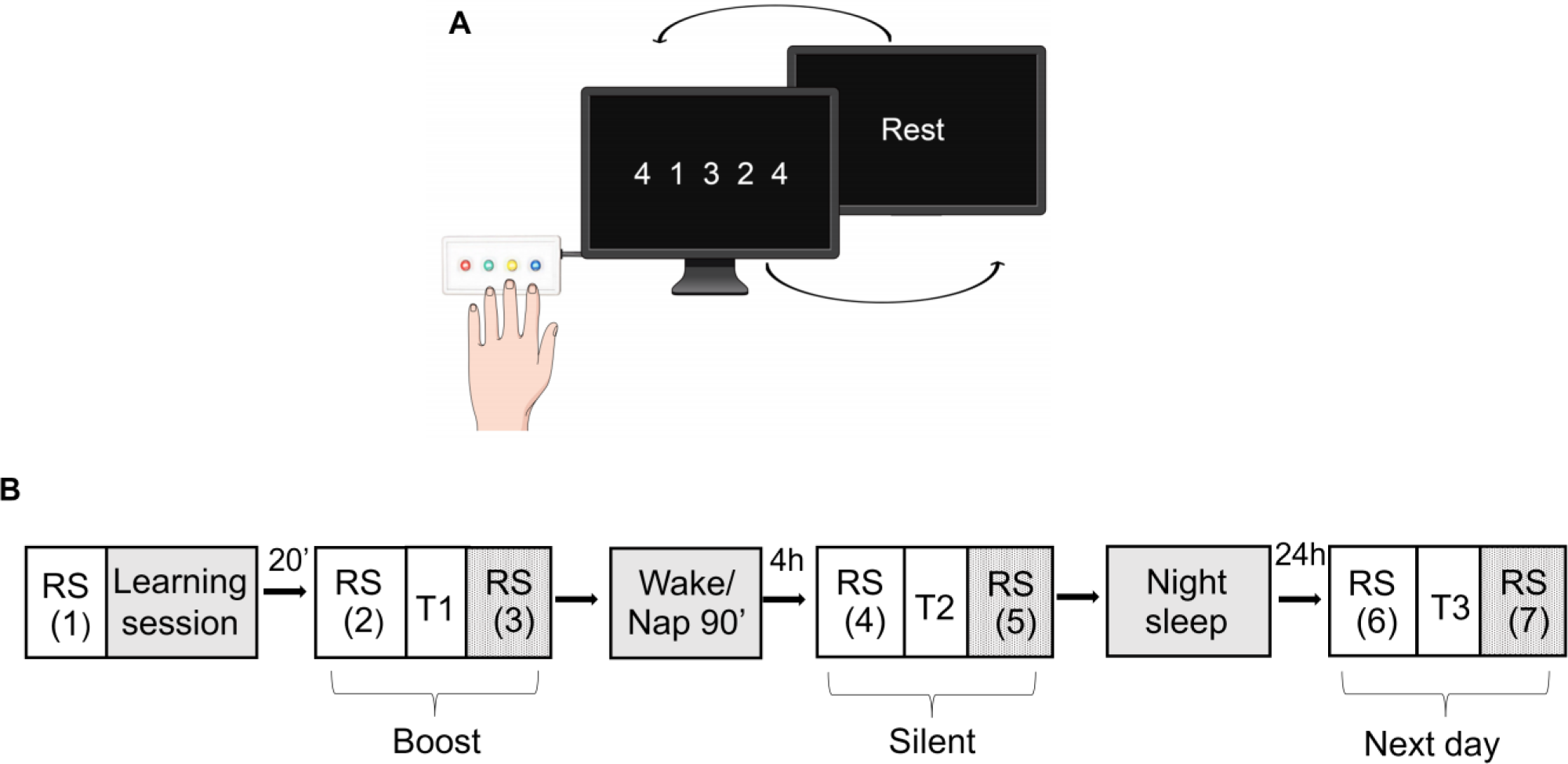
Study design. **A.** One block of the Finger tapping task (FTT): participants perform 20 blocks (30 seconds/block) separated by 20-sec rest intervals on the finger tapping task (FTT) using the four fingers of the left hand (repeated sequence: little/4/, index /1/, ring /3/, middle /2/, little /4/ fingers). **B.** Full experimental design. After a first resting state (RS) recording and a learning session (20 FTT blocks), there were 3 testing windows in the MEG scanner (T1: Boost, T2: Silent, T3: Next day), with each time a RS immediately before and after a short (2 FTT blocks) behavioural testing.

### Behavioural measures of motor performance and learning

For each FTT block, motor performance was estimated by computing the Global Performance Index (GPI), which considers both speed and accuracy [48]. During the learning session, the Best Motor Performance (BMP) was computed by averaging two blocks with the highest GPI scores for each subject. Additionally, we calculated baseline performance averaging the 2^nd^ and 3^rd^ LS blocks (block 1 was excluded since, due to initial habituation to the task, variability was high with frequent stops and disruptions in the execution of the sequence). We also computed a Learning Index (LI) as the percentage change in performance from baseline to BMP. Offline changes in performance were computed from the end of the LS to the first (i.e., boost window), second (silent window), and third (next day window) test sessions (i.e., as percentage change from BMP to best GPI score in the corresponding post-learning test).

### Neuroimaging data acquisition

MEG data acquisition was performed using a 306-channel whole-scalp MEG system (Triux, MEGIN, Helsinki, Finland) at a 1 kHz sampling rate inside a lightweight magnetically shielded room (Maxshield, MEGIN, Helsinki, Finland) at the CUB Hôpital Erasme (Brussels, Belgium). Four coils continuously tracked the subjects’ head position inside the MEG helmet. Coils’ position and about 300 head points were determined following anatomical fiducials with an electromagnetic tracker (Fastrak, Polhemus, Colchester, Vermont, USA). An online analogue band-pass filter was applied for all recordings in the 0.1–330 Hz range.

Nocturnal sleep and naps were recorded at a 1 kHz sampling rate using a MEG-compatible 256-channel scalp EEG system with silver chloride-plated carbon-fibre electrode pellets (MicroCel Geodesic Sensor Net with Net Amp GES 400, Electrical Geodesics Inc., Magstim EGI, Eugene, Oregon, USA). The reference electrode was positioned at Cz, and all impedances were kept below 50 kΩ.

High-resolution 3D T1-weighted cerebral magnetic resonance images (MRIs) were acquired after the last MEG recording on a 1.5 T MRI scanner (Intera, Philips, The Netherlands).

### MEG data pre-processing

The offline temporal signal space separation method [44] was applied to the continuous MEG data to minimise external magnetic interference and head movement corrections (Maxfilter v2.1, MEGIN, Finland). Next, data were filtered (offline band-pass filter: 0.1–45 Hz), and an independent component analysis (FastICA algorithm with dimension reduction to 30 components, hyperbolic tangent nonlinearity function) [48] was applied for visual inspection. Independent components corresponding to cardiac, ocular, and system artifacts were rejected by regressing their time course from the full-rank data. To proceed with source reconstruction, the MEG forward models were estimated based on the participants’ 3D T1-weighted cerebral MRI, anatomically segmented using FreeSurfer software (version 6.0; Martinos Center for Biomedical Imaging, Massachusetts, USA). The MEG and MRI coordinate systems were co-registered via three anatomical fiducials points (nasion and auriculars) for primary head position estimation and the head-surface points for manual refinement (MRIlab, MEGIN Data Analysis Package 3.4.4, MEGIN, Helsinki, Finland). A volumetric and regular 5-mm source grid was constructed in the Montreal Neurological Institute (MNI) template MRI and non-linearly deformed onto each participant’s MRI using the Statistical Parametric Mapping Software (SPM12, Wellcome Centre for Neuroimaging, London, UK). Finally, the three-dimensional MEG forward model associated with this source space was estimated using a one-layer Boundary Element Method executed in the MNE-C suite (Martinos Center for Biomedical Imaging, Massachusetts, USA).

Cleaned pre-processed MEG data were then filtered in one wide frequency band (4–30 Hz). Source activity was reconstructed using MNE [49], which we used here instead of the Beamformer as MNE allows the investigation of RS network connectivity and state dynamics related to the DMN and particularly involving posterior midline cortices (i.e., the precuneus and the posterior cingulate cortex, which are known to be involved in attention, memory and motor-related processes [50]–[53]). The noise covariance matrix was estimated from 5-min empty room MEG recordings spatially filtered using the signal space separation method [44] and temporally filtered between 0.1 and 45 Hz. The MNE regularisation parameter was fixed using the consistency condition derived in [54]. Three-dimensional dipole time courses were projected on their direction of maximum variance, and their Hilbert envelope signal was extracted using the Hilbert transform. These source signals were used both for HMM state inference and for functional connectivity analyses.

### Hidden Markov model (HMM) dynamic analysis

The HMM analysis follows the pipeline described in [34], [55] and is implemented in GLEAN (https://github.com/OHBA-analysis/GLEAN). The number of transient states was set to 8 for consistency with previous MEG power envelope HMM studies [34], [38], [55]–[57]. The 8-state HMM was inferred from the wide-band filtered (4–30 Hz) source envelope signals. Envelope data were downsampled at 10 Hz with a moving-window average with 75% overlap (100 ms wide windows, sliding every 25 ms; resulting in an effective downsampling at 40 Hz), demeaned, normalised by the global variance, and temporally concatenated across participants to design a group-level HMM analysis and across the 7 RS sessions to identify network states common to both the pre- and post-learning sessions (for further discussion on this strategy, see, e.g., [38]). The concatenated envelopes were then pre-whitened and reduced to 40 principal components. Finally, the HMM algorithm [58], [59] was repeatedly run on this dataset 10 times (to account for different initial parameters and retain the model with the lowest free energy) to infer states classifying different power envelope covariance patterns. The Viterbi algorithm was used to decode the binary signals of temporally exclusive state activation/inactivation. Based on these signals, four state temporal parameters were estimated: MLT (mean duration of time intervals of active state), FO (entire fraction of time of the active state), MIL (mean duration of time intervals of inactive state), and NO (total number of state visits). These indices were estimated separately for each state, subject and RS session by deconcatenating the state activation time series. State power maps were obtained as a result of the partial correlation between HMM state activation/inactivation time series and the concatenated source envelope signals, which assess state-specific power changes upon each state activation.

### Statistical contrasts and correlation analyses with HMM state temporal parameters

The comparison among HMM states’ temporal parameters was conducted using repeated measures ANOVA with session (boost, silent, and next day windows; see Fig. 1) and induction (pre vs post-FTT tests) as within-subject factors and group (Wake and Nap) as between-subject factor. Significance was set to *p* < .05 Bonferroni corrected for multiple comparisons by a factor of 21 (7 independent HMM states given that state activation of one state may be predicted from the activation of the 7 other states × 3 independent temporal parameters given that, e.g., NO is strongly dependent on MLT, MIL and FO). Next, we used Spearman’s rank correlation analyses to investigate the relationship between each HMM state’s temporal parameter and behavioural indices. Non-parametric tests were favoured due to higher robustness against outliers, which sometimes arise among HMM state temporal parameters when, e.g., one or a few subjects scarcely visit one state. The calculations were performed using JASP version 0.16.2, JASP Team (2022).

### Resting-state functional connectivity

Source envelope connectivity analysis was performed across a functional connectome that included 126 regions of interest (116 nodes correspond to the Automated Anatomical Labelling (AAL) atlas [60] (including cerebellum), plus 10 nodes based on the relevant motor learning literature) (see Supplementary Table 5; MNI coordinates from the literature were derived/adjusted using SPM Anatomy toolbox [61], Version 2.2b). The resulting 126 × 126 rsFC connectome matrices were computed using the slow amplitude envelope correlation [62] between each node signal and the others, corrected beforehand for spatial leakage using the geometric correction scheme, which prevents false spurious connectivity from dominating over physiological couplings [54]. Additionally, we computed a 126 × 1 vector of power estimate (i.e., source signals’ variance) at each node to control for possible power-induced effects in rsFC changes.

### Statistical analysis with Network Based Statistics

To characterise the functional brain networks engaged in motor learning and consolidation, we performed a statistical within-group comparison of inter-regional connectivity differences using nonparametric NBS [63], [64] on the wideband data (4–30 Hz, same range as the HMM frequency band). We started with mass univariate t-tests looking for learning (pre-vs. post-leaning (RS 1 vs. RS 2)) and induction (pre-vs. post-test RS; e.g., RS 2 vs. RS 3) effects on rsFC connectome matrices, separately within each group. NBS controls the Family-Wise Error Rate (FWER) by identifying ‘network components’ (i.e., adjoining sets of inter-regional connections associated with t values above a pre-defined threshold; here, t > 3.5) whose size is significantly larger than expected by chance. FWER-corrected *p*-values for these network components were generated using n = 5000 random permutations. Furthermore, we conducted a comprehensive correlation analysis to investigate the relationship between connectivity measures and behavioural indices such as Best Motor Performance (BMP) and Learning Index (LI), along with HMM temporal parameters. NBS was again used, but now with FWER-corrected *p*-values for network components generated using Pearson’s correlation and random reshufflings of subjects within behavioural or HMM parameters. For the sake of completeness, we additionally performed separate statistical within-group comparisons of inter-regional connectivity differences using NBS on band-limited rsFC computed within 5 distinct frequency bands: delta (δ 1–4 Hz), theta (θ band: 4–8 Hz), alpha (α band: 8–13 Hz), beta (β band: 13–30 Hz) and gamma (γ band: 30–45 Hz), see Supplementary material.

### State network connectivity strength

To further interpret and connect possible results on HMM and rsFC parameters, we investigated connectivity within ‘state networks’ associated with HMM states showing statistically significant effects. The state network of an HMM state was defined as the subset of nodes of our connectome where the corresponding HMM state power map exceeded a threshold of 60% of its maximum absolute value. State networks for HMM States 3, 7, and 8 (see results section) had 13, 18, and 16 nodes, respectively, and led to rsFC matrices of size 13 × 13, 18 × 18, and 16 × 16, respectively. To quantify the overall connectivity strength of each state network, we first calculated the average of all power envelope correlation values for each node within that specific network, yielding a single value for each node. Then, we obtained the mean of these values across all nodes within each network. This resulted in a single value representing the mean connectivity strength for each subject in each resting-state session, which was used in a subsequent correlational analysis with behavioural measures using JASP version 0.16.2, JASP Team (2022).

### HD-EEG data sleep scoring

Polysomnographic data were downsampled at 250 Hz and converted into European Data Format (EDF). Sleep scoring was performed using PRANA (Polygraphic Recording Analyzer, version 16.01.2007, PhiTools, Strasbourg, France) on a standard montage (10/20 system) with scalp electrodes F3, C3, and O1 referenced to the right mastoid, and F4, C4, and O2 referenced to the contralateral (left) mastoid. Facial electrodes derived electromyographic (EMG) and electrooculographic (EOG) activity. All naps and night recordings were scored using 20-sec epochs with bandpass 0.3–30 Hz for the scalp electrodes, 0.15–15 Hz for EOG channels, and 10–100 Hz for EMG channels. We retained the night scores of 28 participants for analysis as one dataset was lost due to a backup system error, and another one had a corrupted signal.

## RESULTS

### Sleepiness and fatigue measures

A session (before LS, boost, silent and next day sessions) by group (Nap vs. Wake) repeated measures ANOVA was conducted separately on VAS for sleepiness and fatigue. There was a main session effect both for sleepiness (F(2.21, 61.82) = 4.25, *p* = .02) and fatigue (F(3, 84) = 4.48, *p* = .006), respectively. Interaction effects (session × group) were non-significant (sleepiness F(2.21, 61.82) = 2.37, *p* = .10; fatigue F(3, 84) = 0.91, *p* = .44). The results showed a group effect of tiredness F(1, 28) = 4.42, *p* = .04 but not for sleepiness F(1, 28) = 3.87, *p* = .06. The post-hoc analysis revealed that the level of tiredness was significantly higher in the Wake group before testing in the silent window (t(28) = 2.27, *p* = .03).

### Sleep measures

Sleep scoring information for the nap and overnight sleep can be found in Supplementary Information material (Tables 1 and 2, respectively). All participants slept for an average duration of 7.2 ± 0.7 hours during the night. To investigate potential group differences in the duration of sleep stages during the night, we computed independent t-tests for sleep stages (Wake, NREM 1, NREM 2, SWS, REM) and total sleep time (TST) measures. The results did not reveal any significant difference in nocturnal sleep between the Nap and Wake condition groups (all *p*-values ≥ .10).

**Table 1.** Correlations between Best motor performance (BMP) and HMM temporal parameters at pre- and post-learning resting-state sessions for State 3 (Temporal/Sensorimotor) and 7 (Frontal/Cuneus). Spearman’s rho and *p*-values (\*\**p* ≤ 0.002 corrected by factor 21 [7 HMM states × 3 HMM temporal parameters]; \**p* ≤ .017, corrected for 3 HMM temporal parameters).

| HMM temporal parameter | Rest session | RS 1 | RS 2 | RS 3 | RS 4 | RS 5 | RS 6 | RS7 |
| --- | --- | --- | --- | --- | --- | --- | --- | --- |
|  |  | BL | Boost |  | Silent |  | Next day |  |
| **State 3** |  |  |  |  |  |  |  |  |
| FO | Spearman's rho | 0.49 | 0.44 | 0.39 | 0.42 | 0.33 | 0.42 | 0.38 |
|  | *p*-value | **0.006\*** | **0.015\*** | 0.034 | 0.021 | 0.076 | 0.022 | 0.04 |
| MLT | Spearman's rho | 0.47 | 0.47 | 0.47 | 0.39 | 0.39 | 0.37 | 0.32 |
|  | *p*-value | **0.009\*** | **0.008\*** | **0.008\*** | 0.034 | 0.032 | 0.047 | 0.088 |
| MIL | Spearman's rho | -0.44 | -0.14 | -0.37 | -0.35 | -0.35 | 0.31 | 0.38 |
|  | *p*-value | **0.015\*** | 0.449 | 0.043 | 0.061 | 0.055 | 0.102 | 0.039 |
| NO | Spearman's rho | 0.47 | 0.41 | 0.37 | 0.37 | 0.31 | 0.44 | 0.4 |
|  | *p*-value | **0.01\*** | 0.023 | 0.047 | 0.045 | 0.093 | **0.014\*** | 0.028 |
| **State 7** |  |  |  |  |  |  |  |  |
| FO | Spearman's rho | -0.31 | -0.32 | -0.34 | -0.36 | -0.47 | -0.32 | 0.28 |
|  | *p*-value | 0.095 | 0.09 | 0.066 | 0.051 | **0.008\*** | 0.086 | 0.13 |
| MLT | Spearman's rho | -0.45 | -0.42 | -0.29 | -0.21 | -0.39 | -0.27 | 0.14 |
|  | *p*-value | **0.013\*** | 0.021 | 0.115 | 0.271 | 0.033 | 0.148 | 0.446 |
| MIL | Spearman's rho | 0.30 | 0.40 | 0.3 | 0.41 | 0.55 | 0.27 | 0.31 |
|  | *p*-value | 0.105 | 0.028 | 0.108 | 0.025 | **0.002\*\*** | 0.157 | 0.094 |
| NO | Spearman's rho | -0.30 | -0.37 | -0.37 | -0.36 | -0.47 | -0.31 | -0.28 |
|  | *p*-value | 0.109 | 0.041 | 0.046 | 0.052 | **0.009\*** | 0.101 | 0.133 |

**Table 2.** Correlations between Learning Index (LI) and HMM temporal parameters at pre- and post-learning resting-state sessions for State 8 (Cuneus/Somatosensory). Spearman’s Rho and *p*-values (\*\**p* ≤ 0.002; corrected by factor 21 (7 HMM states × 3 HMM temporal parameters); \**p* ≤ .017, corrected for 3 HMM temporal parameters).

| HMM temporal parameter | Rest session | RS 1 | RS 2 | RS 3 | RS 4 | RS 5 | RS 6 | RS7 |
| --- | --- | --- | --- | --- | --- | --- | --- | --- |
|  |  | BL | Boost |  | Silent |  | Next day |  |
| State 8 |  |  |  |  |  |  |  |  |
| FO | Spearman's rho | -0.35 | -0.57 | -0.56 | -0.48 | -0.55 | -0.31 | -0.49 |
|  | *p*-value | 0.058 | **0.001\*\*** | **0.002\*\*** | **0.008\*** | **0.002\*\*** | 0.101 | **0.007\*** |
| MLT | Spearman's rho | -0.33 | -0.43 | -0.4 | -0.46 | -0.56 | -0.28 | -0.52 |
|  | *p*-value | 0.079 | 0.018 | 0.028 | **0.011\*** | **0.002\*\*** | 0.134 | **0.004\*** |
| MIL | Spearman's rho | 0.34 | 0.47 | 0.48 | 0.38 | 0.43 | 0.14 | 0.27 |
|  | *p*-value | 0.064 | **0.009\*** | **0.008\*** | 0.039 | 0.02 | 0.477 | 0.15 |
| NO | Spearman's rho | -0.29 | -0.52 | -0.47 | -0.43 | -0.43 | -0.27 | -0.4 |
|  | *p*-value | 0.119 | **0.003\*** | **0.01\*** | 0.019 | **0.017\*** | 0.146 | 0.028 |

### Motor learning performance

During the learning session, FTT performance (GPI index) rapidly improved and reached asymptotic levels at the end of practice (blocks 1-20), then increased in the boost window (blocks 21-22) to go back at the end of learning level in the silent window (blocks 23-24) and again increased on the next day window (blocks 25-26; Figure 2 A).

**Figure 2.**
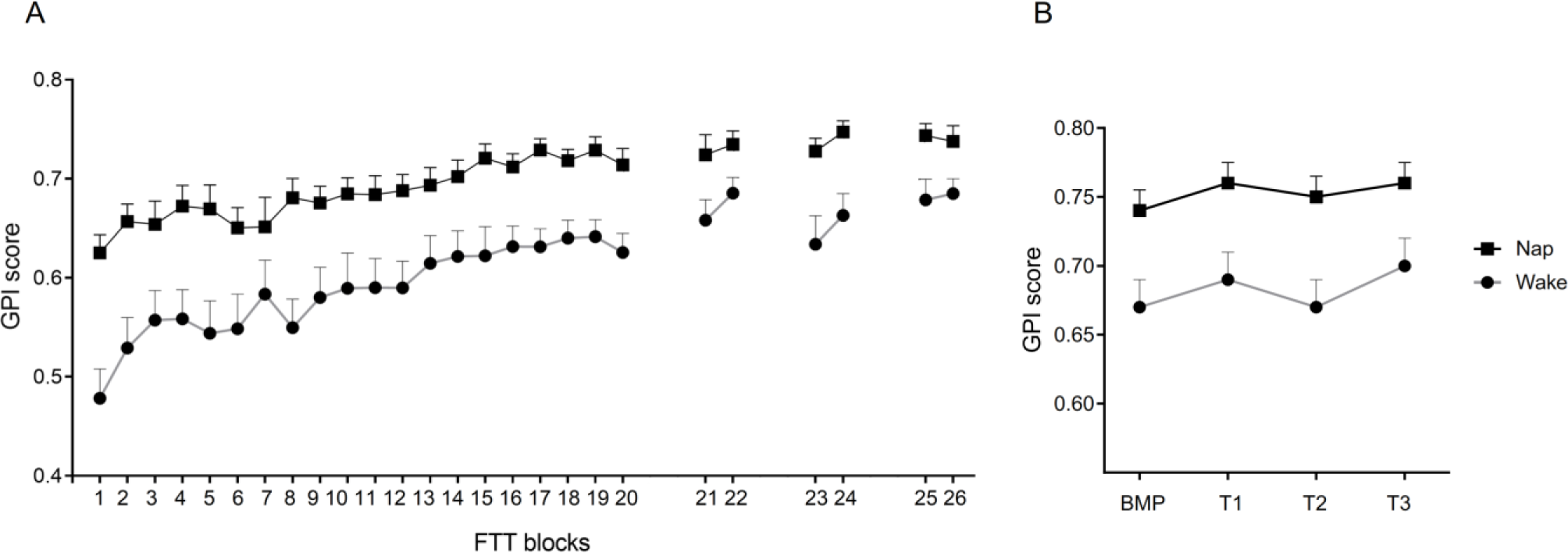
Global Performance Index (GPI) evolution. **A.** Mean GPI scores across 26 blocks of Finger Tapping Task (FTT); blocks 1-20 correspond to the Learning session; blocks 21-22 – boost (T1); blocks 23-24 – silent (T2) and blocks 25-26 – the next day (T3) windows. **B.** GPI evolution from Best Motor Performance (BMP) at the end of the Learning session to each testing session. Circles: Wake condition; Squares: Nap condition. Error bars represent standard errors.

A repeated measures ANOVA computed on GPI scores with the session (BMP × Test 1 × Test 2 × Test 3) and group (Nap vs. Wake) factors disclosed a main session (F(3,84) = 5.45, *p* = .002) and group (F(1,28) = 12.74, *p* = .001) effects. The interaction effect between group and session was non-significant (F(3,84) = 0.78, *p* = .51; Figure 2 B). Post-hoc analyses disclosed performance gains in the boost (BMP vs. T1: *t*(29) = -3.72, *p* = .001) and next day (BMP vs. T3: *t*(29) = -3.34, *p* = .002) windows. Additionally, performance improved from the silent to the next day window (T2 vs. T3: *t*(29) = -2.06, *p* = .05).

### Fast activation/deactivation dynamics of brain networks (HMM analysis)

#### MEG power activation and deactivation of network states

As a reminder, 8 HMM states representing fast resting-state brain network dynamics were inferred from the power envelope of wideband (4–30 Hz) source-reconstructed MEG RS recordings, temporally concatenated over the 7 RS sessions across all participants. The obtained HMM state power maps disclosed distinctive brain topographies related to resting-state networks (RSNs) (Figure 3). State 1 (Frontal/Sensorimotor) featured a network configuration with simultaneously increased power over the prefrontal cortex and decreased power in the bilateral sensorimotor cortices upon state activation. State 2 (Posterior DMN) showed a decreased power in posterior nodes of the default mode network (DMN), including the left and right angular gyri and the posterior midline cortices. Temporal/Sensorimotor State 3 exhibited anti-correlation between power increase in the left temporal cortex and power decrease in the right somatosensory area. Sensorimotor/Visual State 4 featured similar networks as State 8 (Cuneus/Sensorimotor) but with the opposite activation/deactivation patterns, i.e., increased power bilaterally in the sensorimotor areas and decreased power in the visual cortices. State 5 (Calcarine/Postcentral) exhibited power increase and decline in the calcarine and postcentral gyri, respectively. State 6 (Supramarginal) revealed a network where power bilaterally peaked within the supramarginal gyri. The positive component of the power map for State 7 (Frontal/Cuneus) spanned over the prefrontal cortex, while the negative one was located over the right cuneus.

**Figure 3.**
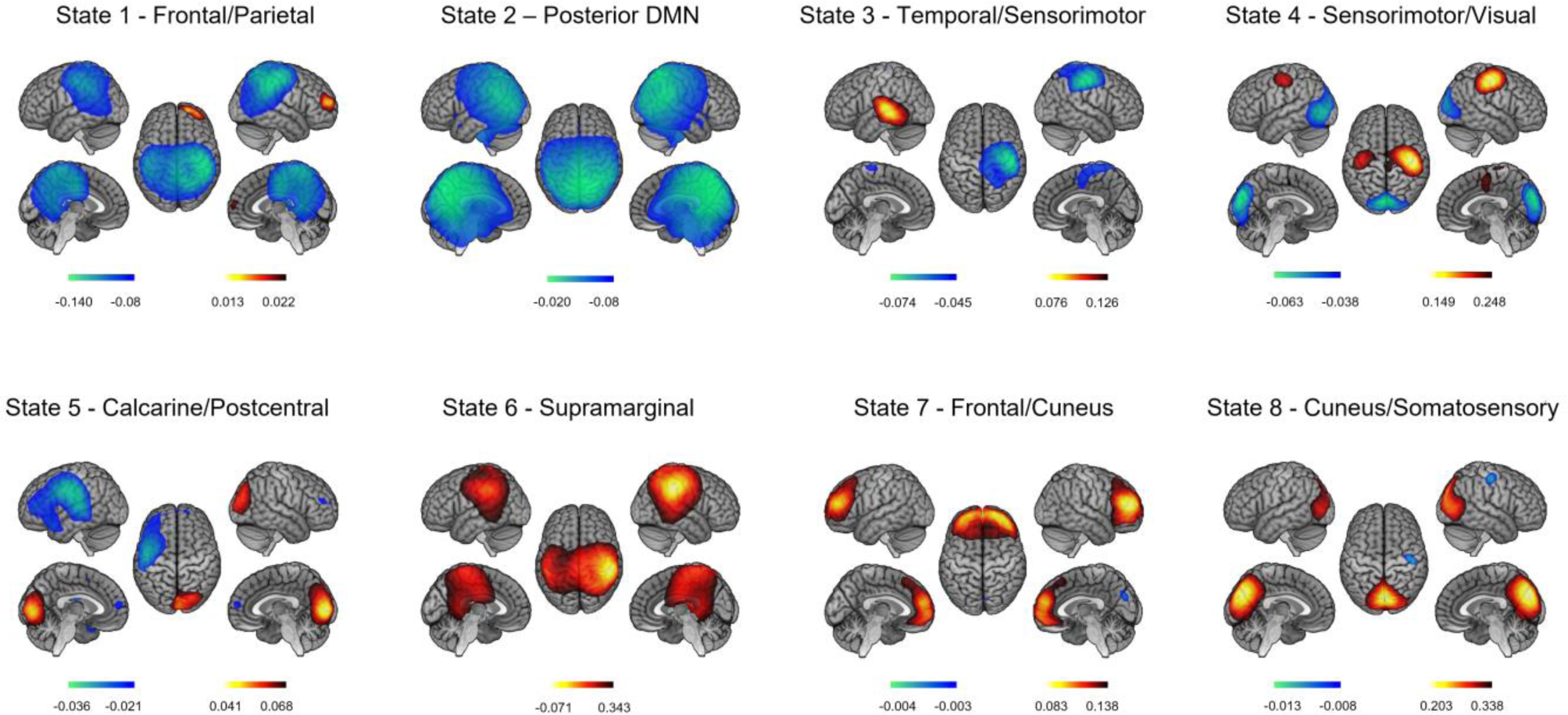
Spatial topographies of HMM transient states computed over the 7 rest sessions (RS1-7). Red/blue scales indicate positive/negative correlation values between the envelope and the state activation/inactivation time course (i.e., increased/decreased power during one state visit). For visualisation purposes, the maps are thresholded to 60% of the maximum absolute of the partial correlation values.

#### HMM temporal characteristics

Separate repeated measures ANOVA with session (T1 vs. T2 vs. T3) and induction (pre– vs. post-behavioural testing) as within-subject factors and group (Nap vs. Wake) as between-subjects factor were computed on the four HMM state temporal parameters (i.e., MLT, mean life time; FO, fractional occupancy; MIL, mean interval length; NO, number of occurrences). The significance level was set at *p* < .05, Bonferroni corrected by a factor of 21 (i.e., 7 independent HMM states and 3 independent HMM temporal parameters).

Results disclosed significant session and induction effects in State 4 (Figure 4). Main session effects were disclosed for FO (F(2,56) = 7.63, *p* = .001), MLT (F(2,56) = 7.60, *p* = .001) and NO (F(2,56) = 8.25, *p* < .001) parameters. There was a trend for MIL (F(2,56) = 2.80, *p* =.07 n.s.). We found a main induction effect in FO (F(1,28) = 5.08, *p* = .032), MIL (F(1,28) = 4.76, *p* = .038) and NO (F(1,28) = 4.52, *p* = .042). The results did not pass the statistical threshold for MLT (F(1,28) = 2.17, *p* = .153). Additionally, we observed a significant interaction effect between session and induction for FO (F(2,56) = 5.46, *p* = .007), MLT (F(2,56) = 4.42, *p* = .02), MIL (F(2,56) = 3.61, *p* = .034) and NO (F(2,56) = 4.48, *p* = .016). Group and session by group effects for State 4 were non-significant for all HMM temporal parameters (all *p*-values > .28; Supplementary Table 3), suggesting a lack of effect of the intermediate nap on fast NDs. Session effects were also observed for HMM states 3, 5 and 6 but did not survive corrections for multiple comparisons (see Supplementary Figures 1-3 and Table 3). Analyses conducted on all other HMM states were non-significant (all *p* - values ≥ .07).

**Figure 4.**
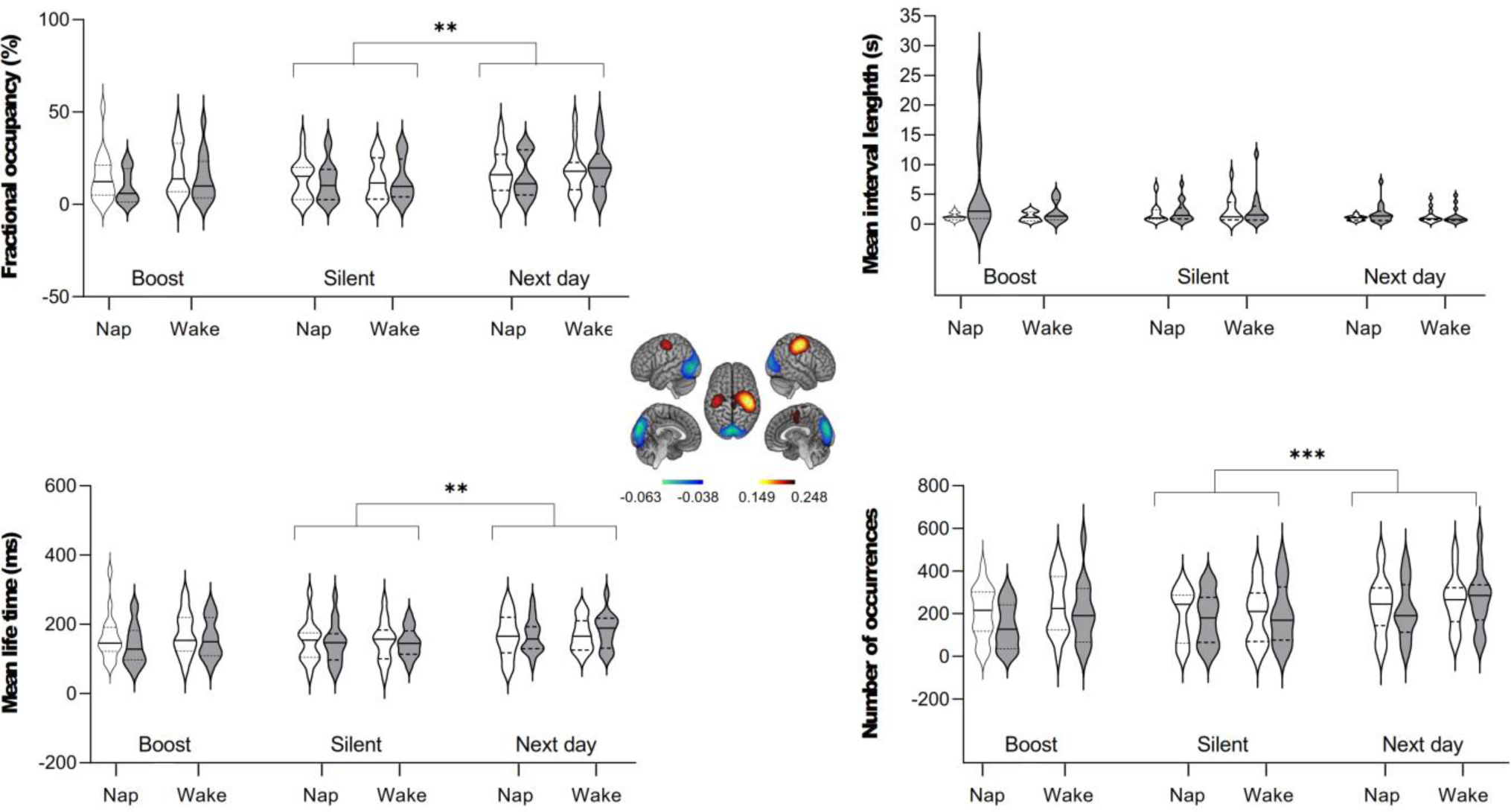
HMM, temporal parameters for Sensorimotor/Visual State 4. White violin plots represent non-induced RS sessions, and grey violin plots induced RS. Medians: solid lines; quartiles: dotted lines. (*** *p* < .001 and ** *p* < .002, corrected by factor 21).

**Table 3.** Correlations between Best Motor Performance (BMP) and state network (corresponding to HMM State 7 (Frontal/Cuneus)). Spearman’s rho and *p*-values; \**p* ≤ .017, corrected by factor 3 (three state networks tested); ^#^*p* ≤ .05, not corrected).

| Rest session | Spearman's rho | p-value |
| --- | --- | --- |
| RS 1 | 0.33 | 0.10 n.s. |
| RS 2 | 0.52 | **0.005\*** |
| RS 3 | 0.42 | **0.028<sup>#</sup>** |
| RS 4 | 0.36 | 0.06 n.s. |
| RS 5 | 0.40 | **0.04<sup>#</sup>** |
| RS 6 | 0.37 | 0.06 n.s. |
| RS 7 | 0.56 | **0.002\*** |

Post-hoc analyses conducted on induction effects within each session in State 4 (paired-sampled Wilcoxon signed-rank tests) showed that re-introduction to the task within the boost window led to significant NDs changes in FO (W = 78, *p* < .001), MIL (W = 415, *p* < .001), NO (W = 68, *p* < .001) (corrected by factor 21) and MLT (W = 115, *p* = .015, uncorrected). Nevertheless, the induction effect did not reach significance during the silent and the next day window (*p* > .129). Overall, these findings suggest that a short evaluation motor task significantly impacted the fast NDs, particularly during the boost window, and that this effect did not persist during the silent and next day windows for this specific HMM state.

Similar analyses conducted on other HMM states revealed a significant induction effect during the silent window solely for State 5 MIL (W = 415, *p* < .001, corrected). Additionally, trends for an induction effect were observed predominantly during the boost and subsequent day windows (*p* > .004 uncorrected, Supplementary Table 4).

#### Associations between HMM state characteristics and behavioural performance/learning

To assess the relationships between fast network NDs and performance, we conducted a correlational analysis between the 8 HMM states temporal parameters and behavioural indices. First, we examined correlations with BMP (i.e., the highest motor performance level achieved by an individual during the learning session). Results disclosed significant correlations with HMM States 3 (Temporal/Sensorimotor) and 7 (Frontal/Cuneus) parameters (see Table 1). Most HMM temporal parameters in State 3 positively correlated with BMP, primarily in baseline RS 1 and pre- and post-learning RS in the boost (RS 2-3) windows, suggesting a trait-like relationship with performance. State 7 exhibited negative correlations with baseline RS 1 and pre- and post-learning RS 4-5 in the silent window.

We then examined the relationships between HMM temporal parameters and the LI, reflecting the intra-individual performance evolution within the learning session. State 8 (Cuneus/Somatosensory) was the only HMM state where temporal parameters consistently exhibited negative correlations with LI (see Table 2).

### Resting-state functional connectivity analyses

As a reminder, analyses were computed within the 4-30 Hz frequency band to make it comparable with HMM results. Three participants were excluded from the analysis due to excessive MEG power values identified during the MEG data quality check, resulting in a final sample of 27 participants. Network-based statistics (NBS) [63], [64] results did not reveal a significant difference between pre- and post-learning RS sessions (RS 2 vs. RS 1). However, re-introduction to the task (induction effect) resulted in the emergence of a vast neural network in the boost window (RS 3 vs. RS 2) comprising 39 edges and 28 nodes (*p* = 0.038; Figure 5). The left temporal middle gyrus was the most heavily weighted node in the network (Supplementary Table 6). Additionally, other network hubs were identified in the left hemisphere, including the superior temporal gyrus and insula, as well as in the right hemisphere, specifically in the posterior medial frontal cortex and supplementary motor area. Induction effects were non-significant in the silent (RS 5 vs. RS 4) and next day (RS 7 vs. RS 6) windows (all *p*-values ≥ .17).

**Figure 5.**
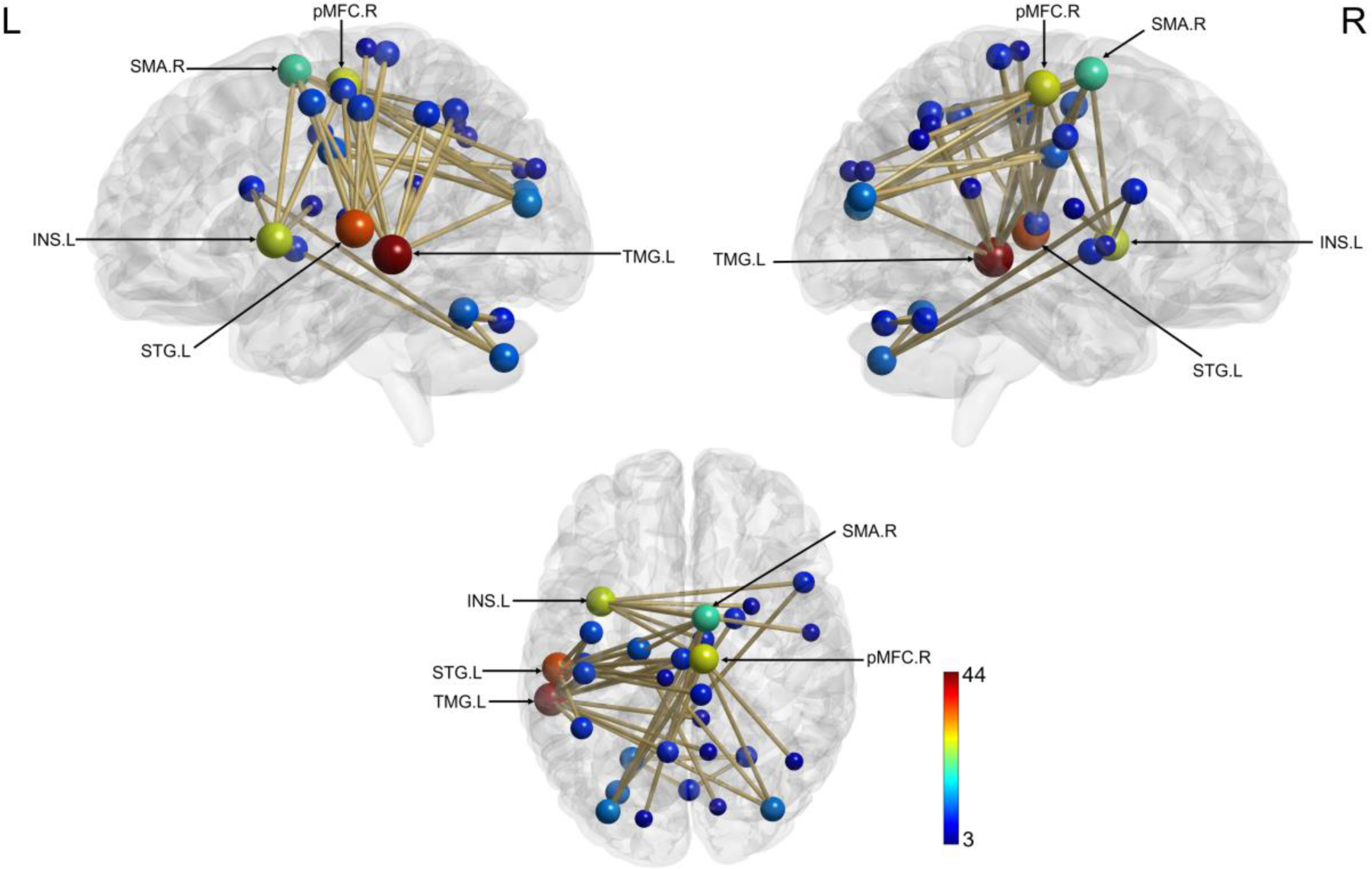
Increased connectivity (NBS analysis) as a result of re-introduction to the task (induction effect) during the boost period. Nodes are scaled according to their weight (the sum of all edges connected to the node). Significant edges are represented as interconnecting lines between 126 connectome seed regions. Abbreviations: TMG – Temporal middle gyrus; STG – Superior temporal gyrus; pMFC - posterior Medial frontal cortex; INS – Insula; SMA – Supplementary motor area; L – left; R – right.

To further investigate changes in rsFC, we conducted a complementary analysis using MEG data filtered in 5 frequency bands. Results disclosed rsFC changes between post- (RS 2) and pre- (RS 1) learning in the theta band (*p* = 0.004) and replicated induction effects in the boost window (RS 3 vs. RS 2) in the delta, alpha, beta and gamma bands (all *p*-values ≤ .03), and in the next day window (RS 7 vs. RS 6) within the alpha and gamma frequency bands (all *p*-values ≤ .05) (Supplementary Figures 4-10). No induction effects were evidenced in the silent window (RS 5 vs. RS 4), all *p* - values ≥ .22.

#### Correlational analysis between connectivity and behavioural indices

Correlational analyses between each RS session and BMP index only highlighted a positive correlation between BMP and RS 3 connectivity (boost window, induced; 29 edges, 24 nodes, *p* = .038; Figure 6). The most weighted node in the correlated network was the right superior frontal gyrus. Additional network hubs were identified in the bilateral middle frontal gyrus and superior frontal gyrus (orbital part) and in the right superior frontal, superior temporal, and angular gyri (Supplementary Table 7).

**Figure 6.**
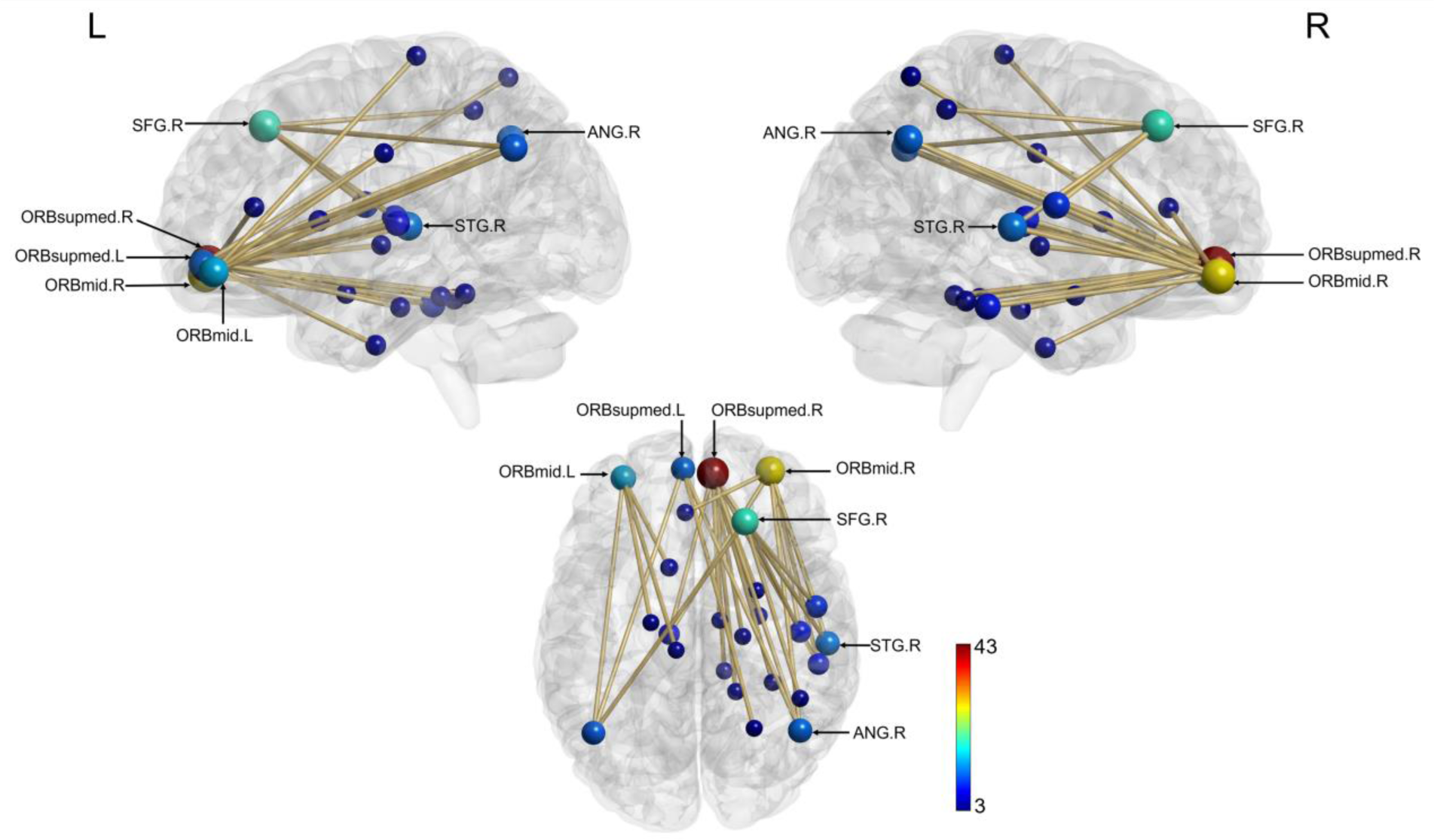
Neural network positively correlated with Best Motor Performance (BMP) Index during RS3 (induced, boost window). Nodes are scaled according to their weight (the sum of all edges connected to the node). Significant edges are represented as interconnecting lines between 126 connectome seed regions. Abbreviations: ORBmid – Middle frontal gyrus, orbital part; ORBsupmed – Superior frontal gyrus, medial orbital; SFG – Superior frontal gurus; ANG – Angular gyrus; STG - Superior temporal gyrus; L-left; R – Right.

As for the other RS sessions, there was a trend towards significance for RS 2 (boost period, non-induced; *p* = .08). All other RS sessions were non-significant (all *p -* values ≥ .09).

Correlational analysis between rsFC and the Learning index revealed significant positive correlations solely for RS 5 (induced RS, silent period) (40 edges, 36 nodes, *p* = .043; Figure 7). The most heavily weighted nodes in the network were located in the right hemisphere, specifically in the thalamus and superior temporal gyrus (Supplementary Table 8). Other network hubs were identified bilaterally in the insula as well as in the left hemisphere in the caudate nucleus and cerebellum crus I, and in the right hemisphere in the hippocampus and fusiform gyrus.

**Figure 7.**
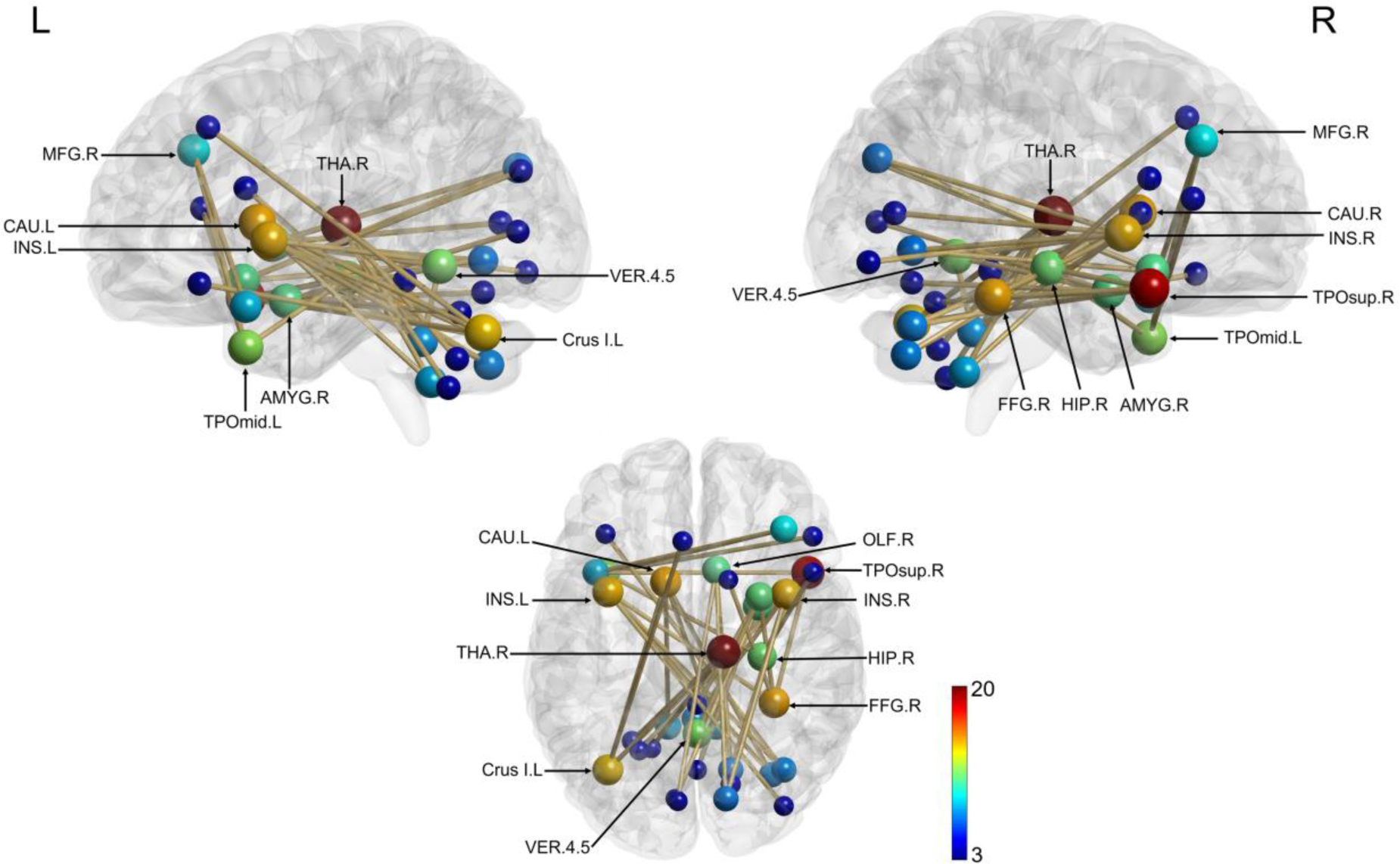
Neural network positively correlated with Learning index during RS5 (induced, silent window). Nodes are scaled according to their weight (the sum of all edges connected to the node). Significant edges are represented as interconnecting lines between 126 connectome seed regions. Abbreviations: MFG – Middle frontal gyrus; CAU – Caudate nucleus; INS – Insula; THA – Thalamus; AMYG – Amygdala; TPOsup – Temporal pole, superior temporal gyrus; TPOmid – Temporal pole, middle temporal gyrus; HIP - Hippocampus; FFG – Fusiform gyrus; Crus I – Crus, Cerebellum; VER – Vermis, Cerebellum; L-left; R – Right.

No significant correlations were disclosed with other RS sessions (all *p*-values ≥ .15).

### State network connectivity and behavioural measures

To examine the associations between HMM power maps, rsFC, and motor performance, we extracted state networks (see Methods) based on the power maps derived by HMM for the 3 states (3, 7 and 8) exhibiting significant correlations with performance. Mean connectivity strength (i.e., the degree of functional connectivity of the whole network, see Methods) for those three HMM state networks was correlated with BMP and LI. Results disclosed positive correlations between BMP and State 7 (Frontal/Cuneus) network (see Supplementary Table 9 for the list of nodes) mean connectivity strength across most of RS sessions (Table 3). Correlation coefficients did not significantly differ between the 7 RS sessions (all *p*-values > .47), suggesting trait-like relationships between HMM parameters and individual motor abilities. The correlations between BMP and mean connectivity strength in the two other state networks did not reach significance (all *p*-values ≥ .18). All correlations with LI were non-significant (all *p*-values ≥ .0.9)

### Relationship between fast and slow neural network dynamics

One of the critical questions we sought to tackle in this study was the relationship between fast (network-level, transient power activation/deactivation) and slow (network connectivity) NDs. To address this question, we performed a correlational analysis between HMM temporal parameters and state networks’ rsFC using the NBS toolbox, focused on the 3 HMM states (States 3, 7, and 8) previously associated with behaviour. Results disclosed negative correlations between connectivity in HMM State 7 (Frontal/Cuneus) with all HMM temporal parameters across the 7 rsFC sessions (Table 4). The number of significant nodes and edges within this state network was highest during the baseline (RS 1) in FO, MLT and NO and decreased to a minimum the next day (RS 7). The correlational outcomes for State 8 did not report any consistent patterns and were below the adjusted statistical threshold (see Supplementary Table 10). Likewise, the analysis for State 3 did not reveal any statistically significant correlations.

**Table 4.**
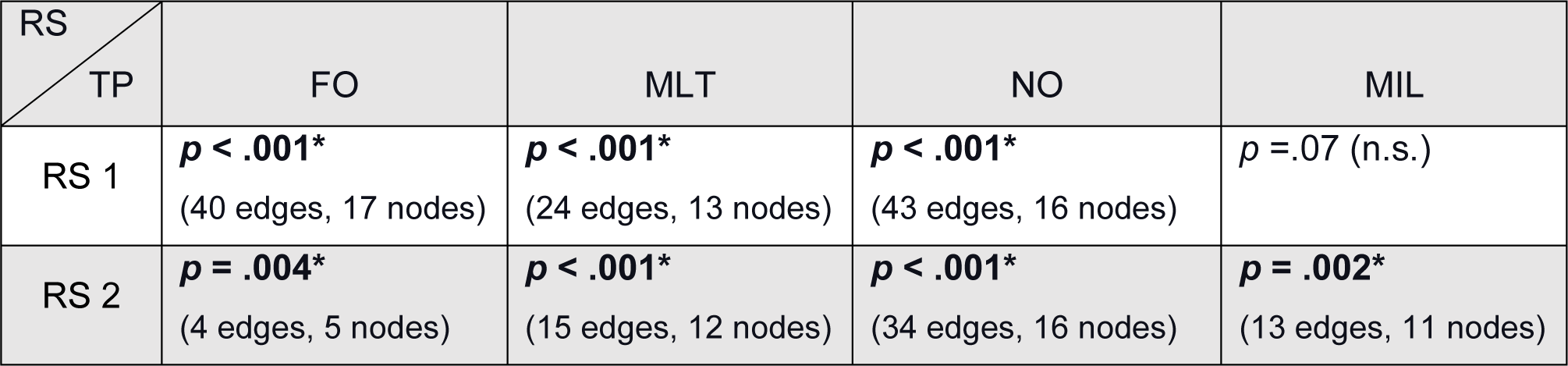
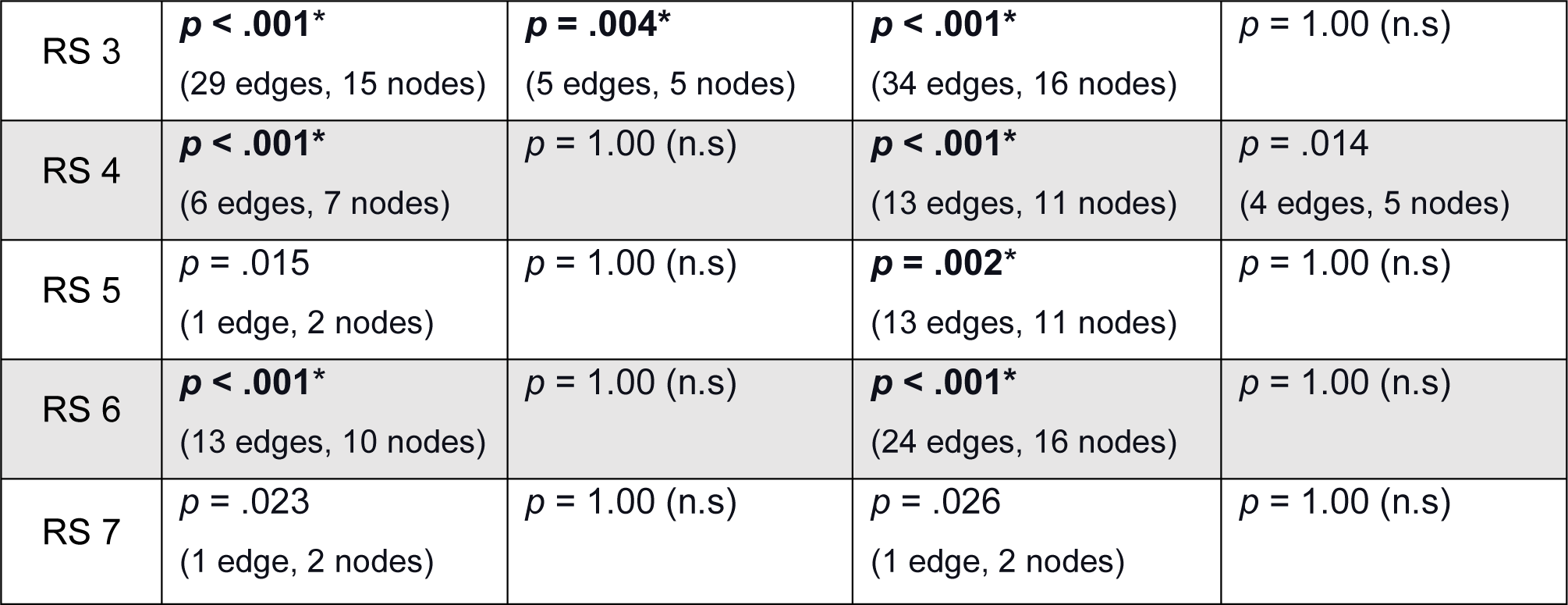
Correlations between HMM temporal parameters (TP) and state network (corresponding to HMM State 7). Spearman’s rho and *p*-values (\**p* ≤ .006, corrected by factor 9; 3 HMM states × 3 HMM TP).

## DISCUSSION

The present study aimed at investigating the neural mechanisms underlying ML during critical learning and consolidation windows, including sleep episodes. We investigated the NDs variations at rest and after the re-introduction to the task (induction effect) within the boost, silent and the next day windows to highlight ML-specific neural networks. We applied both Hidden Markov modelling and resting-state functional connectivity approaches to investigate the NDs changes on fast (sub-seconds) and slow (seconds) timescales, respectively.

### Motor learning performance

Behavioural results showed that the learning curve followed a similar pattern to that described in previous research [4], with distinct variations in performance during the boost, silent and next day windows in the two experimental groups. However, our findings did not reveal significant effects of the intermediate nap on motor performance. These results are consistent with a growing body of literature suggesting that daytime naps do not consistently improve motor memory consolidation [65]–[67], unlike their effect on declarative memory. One possible explanation for the lack of effect is that not all participants in the Nap group reached REM sleep, and those who did only had brief episodes of REM (a few minutes). Indeed, previous studies suggested REM sleep plays a role in integrating new motor skills into existing motor programs [68], eventually leading to more efficient and effective motor performance. Furthermore, according to the sequential hypothesis [69]–[71], optimal learning requires the memory trace to be processed first in non-REM and then in REM sleep, which serve complementary functions in memory consolidation. Therefore, a lack of sufficient REM sleep and/or SWS/REM interplay in our sample might explain the absence of performance improvement in the Nap as compared to the Wake group. Nevertheless, performance gains were noted in Wake and Nap groups following nocturnal sleep, which might either reflect time- or sleep-dependent consolidation. Additionally, nocturnal sleep parameters were similar between the two conditions, suggesting that the post-learning nap did not impact the overnight sleep architecture.

### Associations between MEG network states and behavioural measures

The HMM power maps obtained from the seven resting-state sessions in each participant demonstrated remarkable similarity, like in our previous study in which only the two first RS sessions were used to investigate motor learning-related changes [57]. This further indicates the robustness and stability of the HMM states. Robust associations between motor performance levels achieved at learning and the temporal stability of HMM State 3 (Temporal/Somatosensory) demonstrated a stable, trait-like relationship between the spontaneous organisation of fast recurrent brain networks involved in motor learning and motor ability. Specifically, the observed power decrease in sensorimotor regions might reflect efficient motor execution, planning, and the automation of the motor sequence [72]–[74]. On the other hand, the power increase in temporal regions may represent the engagement of cognitive strategies, potentially linked to internal timing or rhythmic mechanisms. Given that the FTT is a sequential and rapid execution task, it might inherently foster rhythmic abilities. This interpretation aligns with neuroimaging research indicating that rhythm perception relies on the interaction between auditory and motor systems [75], [76]. The robust correlation of HMM state 3 with performance level prior to learning suggests that this state could serve as a neurophysiological marker for predicting individual differences in motor abilities.

On the other hand, HMM state 7 (Frontal/Cuneus) exhibited an opposite relationship with motor performance, although less consistently than State 3. We posit here that this association is tied to the frontal lobe’s function in executive processes, including attentional control and working memory [77], [78]. Moreover, the frontal lobe forms a critical network component involved in sequence learning [79]. As the motor sequence becomes more automatised, frontal areas may become less involved, eventually leading to a decreased engagement in this state. Our results revealed negative correlations solely within the silent window. This might suggest that, during this specific time window, the stabilisation of neural networks occurs, which reflects further automation of performance paralleled by the disengagement of the frontal lobe.

Interestingly, participants who exhibited better learning abilities during the Learning session engaged less in State 8 (Cuneus/Sensorimotor). This suggests that those who most effectively mastered the motor sequence may require less engagement in this state to process motor-related information. Better learners might use more efficient neural strategies, and as they become more proficient at the task, the brain optimises its activity by engaging this state less [80].

Altogether, the correlational findings from the three HMM states suggest that the identified patterns of NDs were not influenced by the particular time of day or the number of sessions used to extract them. Instead, these patterns depict a consistent neural dynamic structure, allowing for optimal system settings for motor learning.

One logical question arises as to why HMM State 4 (Sensorimotor/Visual), which showed significant NDs change over time, did not correlate with performance measures. The fact that changes in the temporal parameters of State 4 were observed during the silent window and the next day could suggest that this state might be engaged in a later stage of motor memory consolidation, such as stabilisation or long-term storage, rather than the initial learning phase that was measured during the learning session.

### Leaning-related resting-state functional connectivity

To assess the relationships between the neural networks underlying motor learning and better understand the functional brain mechanisms revealed by the HMM results, we estimated rsFC matrices for a 126-node connectome in the 4–30 Hz frequency band, similar to the HMM analysis. To some extent, surprisingly, we did not find significant differences between pre- (RS 1) and post- (RS 2) learning RS sessions. It cannot be excluded that rsFC changes induced by the motor learning task were too subtle to be evidenced in this wideband analysis due to a lower signal-to-noise ratio (SNR) compared to analyses conducted on more focused, narrow frequency bands. Additionally, rsFC changes could be frequency specific. To investigate this possibility, we conducted the supplementary analysis using five classical narrow frequency bands from delta to gamma. Our results revealed a significant rsFC change from pre- to post-learning RS sessions (RS 2 – RS 1) solely in the theta frequency band. Theta oscillations have been implicated in encoding and retrieval processes, and their local or long-range synchronisation has been associated with successful memory formation [81]–[83]. The fact that rsFC in the theta band was seen only in the boost window suggests neural adaptations associated with the acquisition and early consolidation of new motor skills, further highlighting the critical role of this immediate post-learning period in memory consolidation processes. Additionally, the vast network that emerged at the early consolidation stage included areas involved in motor control (precentral gyrus, cerebellum, vermis and dentate nucleus [84], [85]), sensory integration (supramarginal gurus [86]) and motor memory processing (hippocampus [15]). The exact mechanisms underlying theta-band rsFC changes and their relationship to motor skill acquisition and consolidation require further investigation.

Our wideband analysis results clearly show that re-introduction to the motor task during the boost window results in widespread changes in neural network connectivity, with the left temporal middle gyrus (TMG) being the weightiest node. Previous research showed grey matter increase in TMG after motor practice [87], [88], suggesting active plastic changes. Other hub elements of this network are also known to be involved in motor performance and learning by enabling planning, execution and control of motor actions (supplementary motor area [89]) as well as monitoring performance problems and ensuring adaptation (posterior medial frontal cortex [90]). Moreover, significant task-induced rsFC changes were observed in the additional 5-band analysis. Interestingly, induction effects were not evidenced during the silent window but were present again on the next day in alpha and gamma frequency bands. Consistent task-related induction of rsFC changes in the boost window might suggest a critical role of this short-lived, transient time window for the early steps of consolidation when newly acquired motor memories are particularly susceptible to reactivation by the task. This reinforces the hypothesis that a boost window within 30 minutes after the end of learning represents a time of high neural activity and plasticity. On the other hand, the lack of induced connectivity changes during the silent window suggests that memory traces may have been stabilised and would thus be less labile. On the next day, induction effects were significant but reduced as compared to the boost window, suggesting that memory traces have been solidified (i.e., consolidated). With elapsed time and sleep, memory traces undergo substantial modifications leading to network reorganisation and information transfer to pre-existing networks [71], [91].

### Correlational analysis between NBS and behavioural measures

BMP achieved during learning positively correlated with increased rsFC during the boost window, particularly after task re-activation (RS 3). Connectivity changes might be driven by neural plasticity associated with the re-introduction to the task within a plastic window, which may lead to the formation of new connections and the strengthening of existing ones. It is worth mentioning that there was also a trend for correlation in RS 2 prior to induction, suggesting that connectivity changes in the boost window are related to the early steps in the consolidation of motor memory, irrespectively of induction. Correlations were non-significant during the silent and next day windows, suggesting that the boost window provides specific neural activity and plasticity ([92], [93] conditions that promote patterns of connectivity changes associated with motor memory reorganisation and consolidation [17].

### Fast and slow neural dynamics

In this study, we also aimed at investigating the relationships between fast transient NDs disclosed by HMM analyses and slower neural activity as reflected by functional connectivity (FC). Although HMM analyses highlighted transient networks with spatial topographies close to part of the RS networks, the link between these two measures still needs to be clarified, which might partly stem from core methodological differences. Indeed, whereas HMM is based on power envelope covariance, FC relies on correlation metrics. Notwithstanding, Seedat and colleagues [94] showed that rsFC correlates with burst coincidence, suggesting that transient, short-lived power bursts underlie the functional connectivity of RS networks. However, we found negative correlations between HMM State 7 network’s temporal parameters and rsFC before ML, suggesting an inverse relationship between bursts and connectivity in this state. It also indicates that the associations between HMM temporal parameters and rsFC are not simply due to the influence of motor practice since both measures already correlate at baseline (RS 1) before learning but rather reflect trait-level characteristics of neural processing. When considering relationships between these neural measures and behaviour, the picture becomes more complex. Better motor performance was both related to reduced associations with HMM temporal parameters and increased connectivity. It suggests that motor skills may be supported by more stable, consistent patterns of neural activity than by transient, variable NDs [57]. This hypothesis is supported by the fact that HMM State 3 and 8 networks exhibited robust correlations with either BMP or Learning Index in terms of HMM temporal parameters but not of connectivity measures. Additionally, temporal parameters were not associated with connectivity in these two state networks.

## Conclusion

To sum up, the present study offers insights into the NDs underlying motor learning and the role of sleep in memory consolidation. Fast NDs revealed by HMM states appear to primarily reflect a trait-like neural processing predictive of individual motor abilities. On the other hand, slower NDs identified by rsFC highlight the critical role of the transient boost window for motor learning, with re-introduction to the task eliciting neural dynamic changes most prominently in this period. Induction has no effect in the silent window, suggesting a stabilised, less malleable neural trace. The next day, induction effects were observable again but reduced compared to the boost window, possibly reflecting sleep- or time-dependent re-organisation of ML-related neural networks. Differential associations between performance and HMM and connectivity parameters suggest that motor skills are supported by more stable, consistent patterns of neural activity than by transient, variable NDs. Overall, it highlights the importance of integrating multiple methods to understand NDs in motor learning comprehensively.

## Acknowledgements

This study was supported by the Fonds de la Recherche Scientifique (FRS-FNRS, Brussels, Belgium; Research Convention Excellence of Science EOS MEMODYN #30446199). Lillia Roshchupkina is a ULB Mini-ARC grant holder previously supported by an FRS-FNRS Research Fellow grant. Nicolas Coquelet was supported by EOS MEMODYN. Xavier De Tiège is/was a Postdoctorate Clinical Master Specialist at the FRS-FNRS.

The MEG project at the CUB Hôpital Erasme is financially supported by the Fonds Erasme (Research Convention: “Les Voies du Savoir”, Brussels, Belgium).

## Authors’ contributions

L.R. and P.P. designed the study; L.R., N.C., and V.W. acquired the data; N.C. and V.W. contributed to the analysis tools; L.R., N.C., P.P., C.U., X.D.T. and V.W. analysed the data and wrote/reviewed the manuscript.

