## Supplementary Information for "Motor learning- and consolidation-related resting state fast and slow brain dynamics across wake and sleep"

Liliia Roshchupkina [1-3]*, Vincent Wens [2-4], Nicolas Coquelet [2-4], Charline Urbain [1-3], Xavier de Tiege [2-4], Philippe Peigneux [1,2]

[1] UR2NF - Neuropsychology and Functional Neuroimaging Research Unit affiliated at CRCN - Centre for Research in Cognition and Neurosciences, Université libre de Bruxelles (ULB), Brussels, Belgium.

[2] UNI - ULB Neuroscience Institute, Brussels, Belgium.

[3] LN²T - Laboratoire de Neuroanatomie et Neuroimagerie translationnelles, ULB, Brussels, Belgium.

[4] Department of Functional Neuroimaging, Service of Nuclear Medicine, HUB - Hôpital Universitaire de Bruxelles, Hospital Erasme, Brussels, Belgium.

### 1. Sleep measures

Polysomnographic data (naps and nights) were scored using 20-sec epochs with bandpass 0.3-30 Hz for the scalp electrodes.

**Table 1. Sleep scoring in the 90-min nap conditions**

| Sleep Stage | Nap sleep (min) | Nap sleep (%) |
|---|---|---|
| Wake | 12.31 ± 3.30 (46.67) | 13.59 ± 3.59 (49.08) |
| NREM 1 | 9.88 ± 1.00 (14.66) | 11.01 ± 1.12 (16.30) |
| NREM 2 | 28.17 ± 3.04 (47.33) | 31.37 ± 3.37 (52.99) |
| SWS | 32.98 ± 4.40 (60.33) | 36.85 ± 4.96 (66.77) |
| REM | 6.87 ± 2.33 (16.33) | 12.75 ± 2.68 (26.56) |
| TST | 89.77 ± 0.43 (43.00) | 86.41 ± 3.59 (43.15) |

**Table 1.** Sleep scoring in the 90-min nap conditions. Mean values and standard errors indicate the duration (min) and percentage (%) of time spent in each stage; the range is indicated in brackets. The range of data is in brackets. Abbreviations: NREM (non-rapid eye movement sleep), SWS (slow wave sleep), REM (rapid-eye movement sleep), TST (total sleep time, considers all sleep stages except wakefulness).

**Table 2. Nocturnal sleep scorings in the Wake and Nap groups**

| Sleep Stage | Wake group (min) | Wake group (%) | Nap group (min) | Nap group (%) |
|---|---|---|---|---|
| Wake | 21.83 ± 5.81 (76.66) | 4.75 ± 1.27 (17.01) | 33.53 ± 10.19 (121.33) | 7.26 ± 2.11 (25.29) |
| NREM 1 | 28.67 ± 3.20 (43.00) | 6.21 ± 0.66 (8.37) | 36.71 ± 4.38 (52.66) | 8.34 ± 1.07 (14.57) |
| NREM 2 | 133.86 ± 12.73 (159.00) | 28.88 ± 2.52 (29.83) | 136.05 ± 9.49 (138.00) | 30.83 ± 2.25 (31.05) |
| SWS | 178.10 ± 12.06 (144.67) | 38.96 ± 2.75 (31.00) | 150.43 ± 8.83 (126.33) | 33.75 ± 1.72 (22.36) |
| REM | 97.96 ± 6.22 | 21.19 ± 1.19 | 88.26 ± 4.88 | 19.81 ± 0.97 |

| | | | | |
|---|---|---|---|---|
| | (88.00) | (17.94) | (48.33) | (9.89) |
| TST | 438.58 ± 10.25<br>(91.00) | 95.25 ± 1.27<br>(17.23) | 411.45 ± 10.58<br>(125.67) | 92.74 ± 2.11<br>(25.29) |

**Table 2.** Nocturnal sleep scorings in the Wake and Nap groups. The mean values and standard errors indicate the duration (min) and percentage (%) of time spent in each stage; the range is indicated in brackets. Abbreviations: NREM (non-rapid eye movement sleep), SWS (slow wave sleep), REM (rapid-eye movement sleep), TST (total sleep time, considers all sleep stages except wakefulness).

### 2. HMM temporal characteristics

To reveal the impact of motor learning (ML) on HMM temporal parameters, we performed a repeated measures ANOVA with Sessions (T1 vs. T2 vs. T3) and Induction (pre – vs. post-behavioural testing) as within-subject factors and Group (Nap vs. Wake) as between-subjects factor. The significance level was set at $p < .05$, Bonferroni corrected by a factor of 21 (i.e., 7 independent HMM states and 3 independent HMM temporal parameters).

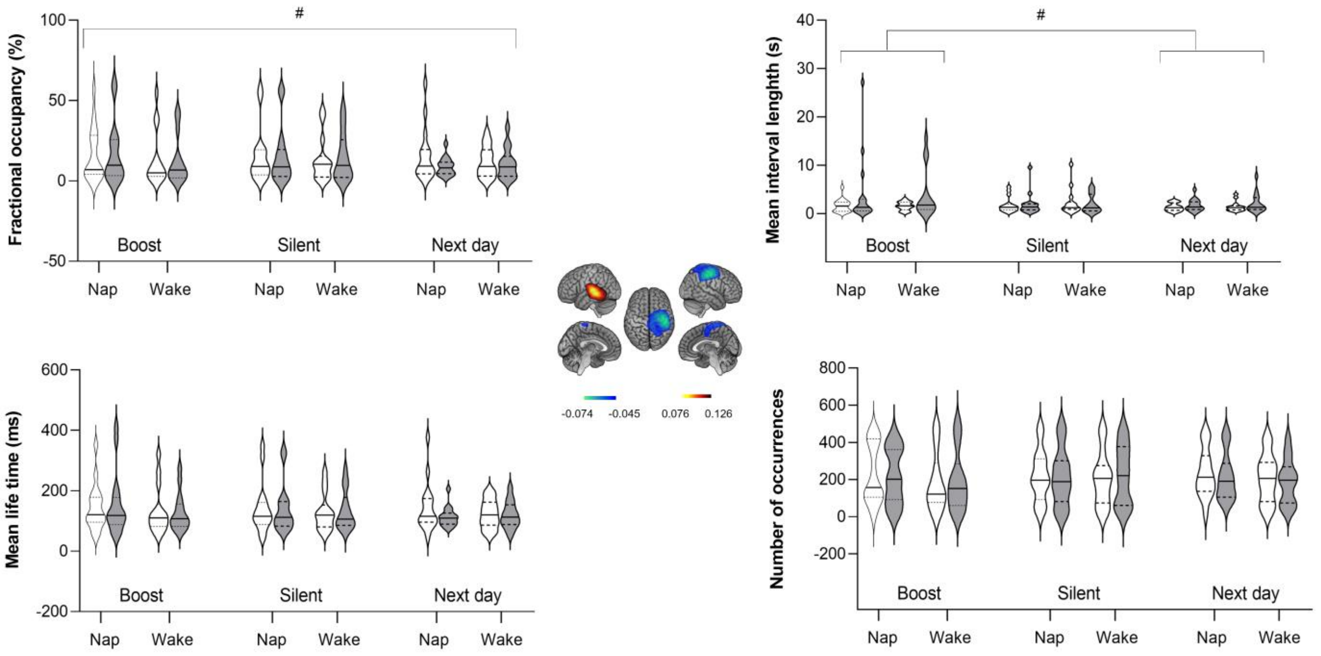


**Figure 1.** HMM temporal parameters for State 3 (Temporal/Sensorimotor). White violins represent non-induced RS sessions, grey violins – represent induced RS, and the medians are in solid lines and quartiles - dotted lines. ( $^{\#}p \leq .05$ not corrected for multiple comparisons).

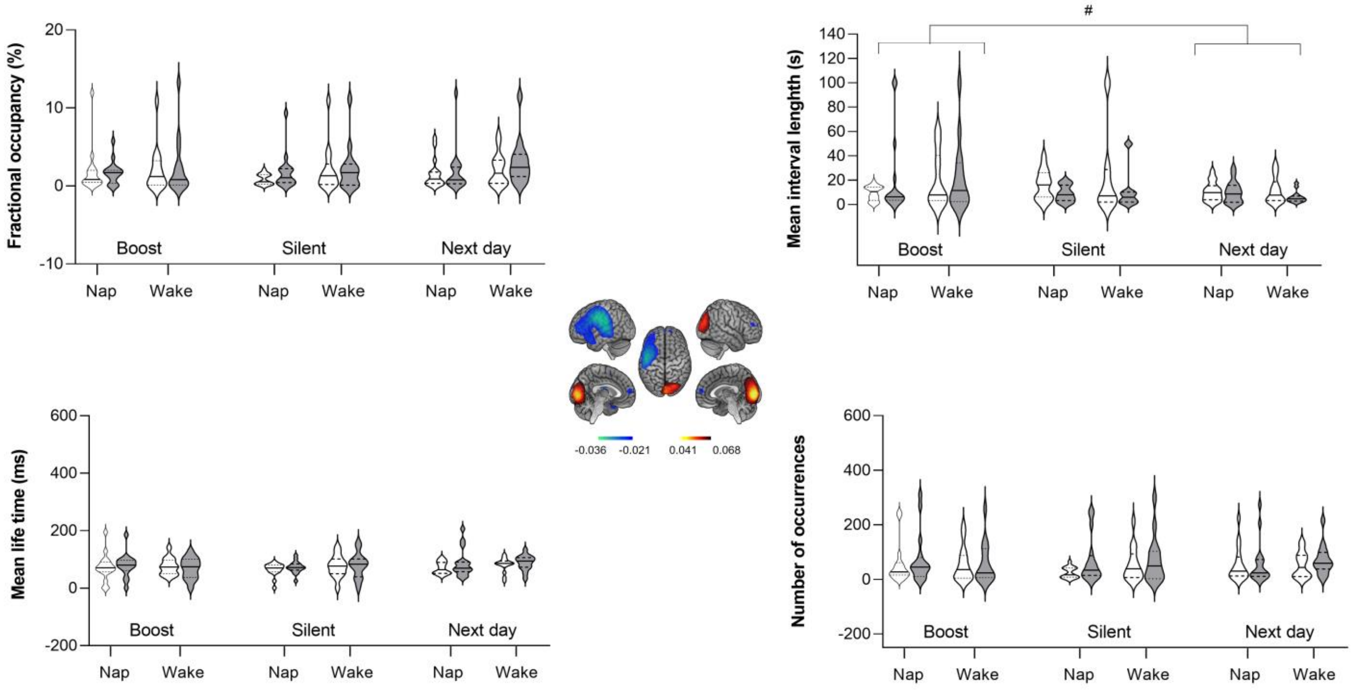


**Figure 2.** HMM temporal parameters for State 5 (Calcarine/Postcentral). White violins represent non-induced RS sessions, grey violins – represent induced RS, and the medians are in solid lines and quartiles - dotted lines. ($^{\#}p \leq .05$ not corrected for multiple comparisons).

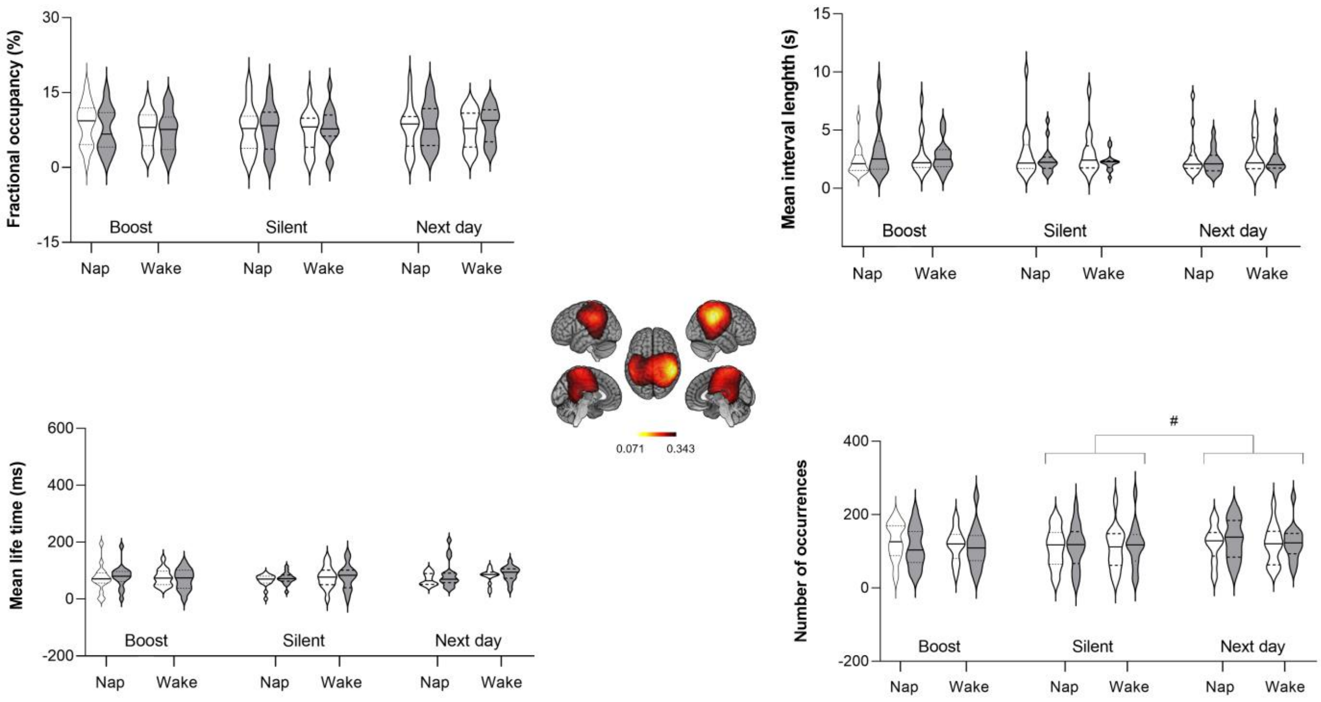


**Figure 3.** HMM temporal parameters for State 6 (Supramarginal). White violins represent non-induced RS sessions, grey violins – represent induced RS, and the medians are in solid lines and quartiles - dotted lines. ($^{\#}p \leq .05$ not corrected for multiple comparisons).

Table 3 represents the results of repeated measures ANOVA for States 3, 4, 5 and 6. Figures 3-4 illustrate the results for these HMM states.

**Table 3. HMM temporal parameters (repeated measure ANOVA) results**

| **Sate 3** | | | | |
|---|---|---|---|---|
| | FO | MLT | NO | MIL |
| Session | F = 3.16<br>***p* = 0.05#** | F = 2.91<br>*p* = 0.06 | F = 0.26<br>*p* = 0.77 | F = 3.54<br>***p* = 0.04#** |
| Session x Group | F = 0.58<br>*p* = 0.56 | F = 0.46<br>*p* = 0.64 | F = 1.15<br>*p* = 0.32 | F = 0.10<br>*p* = 0.91 |
| Induction | F = 2.92<br>*p* = 0.1 | F = 1.73<br>*p* = 0.20 | F = 0.58<br>*p* = 0.45 | F = 4.97<br>***p* = 0.03#** |
| Session x Induction | F = 3.0<br>*p* = 0.06 | F = 1.79<br>*p* = 0.18 | F = 1.26<br>*p* = 0.30 | F = 3.90<br>***p* = 0.03#** |
| Group | F = 0.58<br>*p* = 0.56 | F = 0.23<br>*p* = 0.64 | F = 0.18<br>*p* = 0.68 | F = 0.09<br>*p* = 0.77 |
| **State 4** | | | | |
| Session | F = 7.63<br>***p* = 0.001**** | F = 7.60<br>***p* = 0.001**** | F = 8.25<br>***p* < 0.001***** | F = 2.80<br>*p* = 0.07 |
| Session x Group | F = 1.29<br>*p* = 0.28 | F = 0.60<br>*p* = 0.56 | F = 0.99<br>*p* = 0.38 | F = 0.03<br>*p* = 0.97 |
| Induction | F = 5.08<br>***p* = 0.03#** | F = 2.16<br>*p* = 0.15 | F = 4.52<br>***p* = 0.04#** | F = 4.76<br>***p* = 0.04#** |
| Session x Induction | F = 5.46<br>*p* = **0.007#** | F = 4.22<br>***p* = 0.02#** | F = 4.47<br>***p* = 0.02#** | F = 3.61<br>***p* = 0.03#** |
| Group | F = 0.42<br>*p* = 0.52 | F = 0.20<br>*p* = 0.66 | F = 0.55<br>*p* = 0.46 | F = 0.08<br>*p* = 0.77 |
| **State 5** | | | | |
| Session | F = 0.47<br>*p* = 0.63 | F = 1.11<br>*p* = 0.34 | F = 0.18<br>*p* = 0.84 | F = 4.48<br>***p* = 0.02#** |
| Session x Group | F = 0.13<br>*p* = 0.88 | F = 0.63<br>*p* = 0.54 | F = 0.86<br>*p* = 0.43 | F = 1.08<br>*p* = 0.35 |
| Induction | F = 8.13<br>***p* = 0.008#** | F = 5.60<br>***p* = 0.03#** | F = 4.45<br>***p* = 0.04#** | F = 0.54<br>*p* = 0.47 |
| Session x Induction | F = 0.60<br>*p* = 0.55 | F = 1.63<br>*p* = 0.20 | F = 2.19<br>*p* = 0.12 | F = 4.89<br>***p* = 0.01#** |
| Group | F = 1.08<br>*p* = 0.31 | F = 0.15<br>*p* = 0.70 | F = 0.06<br>*p* = 0.81 | F = 0.47<br>*p* = 0.50 |
| **State 6** | | | | |
| Session | F = 1.34<br>*p* = 0.27 | F = 0.05<br>*p* = 0.96 | F = 3.83<br>***p* = 0.03#** | F = 0.62<br>*p* = 0.54 |
| Session x Group | F = 0.70<br>*p* = 0.50 | F = 1.65<br>*p* = 0.20 | F = 0.27<br>*p* = 0.76 | F = 0.21<br>*p* = 0.81 |
| Induction | F = 0.70<br>*p* = 0.50 | F = 0.35<br>*p* = 0.56 | F = 0.03<br>*p* = 0.86 | F = 1.47<br>*p* = 0.24 |

| Session x Induction | F = 3.82<br>***p* = 0.03#** | F = 2.26<br>*p* = 0.12 | F = 5.20<br>***p* = 0.009#** | F = 3.94<br>***p* = 0.03#** |
|---|---|---|---|---|
| Group | F = 0.03<br>*p* = 0.86 | F = 0.01<br>*p* = 0.91 | F = 0.03<br>*p* = 0.87 | F = 0.01<br>*p* = 0.94 |

**Table 3.** HMM temporal parameters (repeated measure ANOVA) results for States 3, 4, 5 and 6; *p* – values (****p* < 0.001; ***p* ≤ 0.002, corrected by factor 21 (7 HMM states x 3 HMM temporal parameters); #*p* ≤ 0.05, uncorrected).

Table 4 represents the results of post-hoc analyses conducted on Induction effects within each session using a paired-sampled Wilcoxon signed-rank test for all HMM states.

**Table 4. Induction effect for 8 HMM states**

| **Sessions** | **Fractional occupancy** | | **Mean life time** | | **Number of occurrences** | | **Mean interval length** | |
|---|---|---|---|---|---|---|---|---|
| | W | P-values | W | P-values | W | P-values | W | P-values |
| **State 1** | | | | | | | | |
| Boost | 349 | **0.02#** | 349 | **0.02#** | 219 | 0.98 | 159 | 0.14 |
| Silent | 184 | 0.33 | 187 | 0.36 | 138.5 | **0.05#** | 224 | 0.87 |
| Next day | 196.5 | 0.47 | 245 | 0.81 | 104 | **0.01#** | 370 | **0.004#** |
| **State 2** | | | | | | | | |
| Boost | 200 | 0.52 | 242 | 0.86 | 161 | 0.14 | 358 | **0.01#** |
| Silent | 247 | 0.78 | 228 | 0.94 | 263 | 0.33 | 286 | 0.28 |
| Next day | 228 | 0.93 | 207 | 0.61 | 243 | 0.59 | 322 | 0.07 |
| **State 3** | | | | | | | | |
| Boost | 227 | 0.92 | 237 | 0.94 | 198.5 | 0.69 | 281 | 0.33 |
| Silent | 263 | 0.54 | 264 | 0.53 | 230 | 0.55 | 167 | 0.42 |
| Next day | 118 | **0.03#** | 173 | 0.23 | 99 | **0.02#** | 361 | **0.01#** |
| **State 4** | | | | | | | | |
| Boost | 78 | **< .001*****| 115 | **0.02#** | 68 | **< .001***** | 415 | **< .001***** |
| Silent | 168 | 0.19 | 198 | 0.49 | 211 | 0.67 | 322 | 0.07 |
| Next day | 219 | 0.79 | 273 | 0.42 | 175.5 | 0.37 | 307 | 0.13 |
| **State 5** | | | | | | | | |

| | | | | | | | | |
|---|---|---|---|---|---|---|---|---|
| Boost | 193 | 0.67 | 248 | 0.76 | 223 | 0.42 | 252 | 0.70 |
| Silent | 303 | **0.01#** | 290.5 | 0.12 | 283.5 | **0.02*** | 75 | **0.001**** |
| Next day | 313 | 0.10 | 338 | **0.03#** | 309.5 | 0.12 | 123 | **0.03#** |
| **State 6** | | | | | | | | |
| Boost | 152 | 0.16 | 214 | 0.72 | 107.5 | **0.01#** | 353 | **0.01#** |
| Silent | 280 | 0.34 | 276 | 0.38 | 243 | 0.84 | 159 | 0.13 |
| Next day | 325 | 0.06 | 328 | **0.05#** | 266 | 0.30 | 158 | 0.13 |
| **State 7** | | | | | | | | |
| Boost | 329 | **0.05#** | 263 | 0.54 | 245.5 | 0.55 | 200 | 0.52 |
| Silent | 98 | **0.01#** | 158 | 0.20 | 111.5 | **0.02#** | 235 | 0.97 |
| Next day | 110 | **0.02#** | 232 | 1.00 | 107 | **0.01#** | 352 | **0.01#** |
| **State 8** | | | | | | | | |
| Boost | 354 | **0.01#** | 344 | **0.02#** | 289.5 | 0.12 | 93 | **0.004#** |
| Silent | 265 | 0.52 | 261 | 0.57 | 219.5 | 0.97 | 217 | 0.76 |
| Next day | 265 | 0.52 | 318 | 0.08 | 191 | 0.57 | 200 | 0.96 |

**Table 4.** Induction effect for 8 HMM states. W - Wilcoxon signed-rank; (***$p < 0.001$; **$p \leq 0.002$, corrected by factor 21 (7 HMM states x 3 HMM temporal parameters); $^{\#}p \leq 0.05$, uncorrected).

### 3. Resting-state functional connectivity (rsFC) wideband

The functional connectome of our study included 126 regions of interest (116 nodes correspond to the Automated Anatomical Labelling (AAL) atlas [1] (including the cerebellum). Additionally, we added ten nodes based on the relevant learning/memory literature (see Table 5). The MNI coordinates from the literature were derived/adjusted using the SPM Anatomy toolbox, Version 2.2b. To identify the network of regions engaged in motor learning and consolidation, we computed the adjacency matrix (126 by 126) for wideband (4–30 Hz). A statistical within-group comparison of inter-regional connectivity differences was then computed on the adjacency matrix using nonparametric Network Based Statistics (NBS) [2], [3]. Our results reveal that re-introducing the task (induction effect) resulted in a vast neural network in the boost window (RS 3 vs. RS 2) comprising 39 edges and 28 nodes $p = 0.038$. Table 6 represents the nodes list of this emerged network.

### Table 5. Full connectome list of nodes

| Label | x | y | z | Region | Reference |
|---|---|---|---|---|---|
| 1 | -39 | -6 | 51 | Precentral_L | AAL |
| 2 | 41 | -8 | 52 | Precentral_R | AAL |
| 3 | -18 | 35 | 42 | Frontal_Sup_L | AAL |
| 4 | 22 | 31 | 44 | Frontal_Sup_R | AAL |
| 5 | -17 | 47 | -13 | Frontal_Sup_Orb_L | AAL |
| 6 | 18 | 48 | -14 | Frontal_Sup_Orb_R | AAL |
| 7 | -33 | 33 | 35 | Frontal_Mid_L | AAL |
| 8 | 38 | 33 | 34 | Frontal_Mid_R | AAL |
| 9 | -31 | 50 | -10 | Frontal_Mid_Orb_L | AAL |
| 10 | 33 | 53 | -11 | Frontal_Mid_Orb_R | AAL |
| 11 | -48 | 13 | 19 | Frontal_Inf_Oper_L | AAL |
| 12 | 50 | 15 | 21 | Frontal_Inf_Oper_R | AAL |
| 13 | -46 | 30 | 14 | Frontal_Inf_Tri_L | AAL |
| 14 | 50 | 30 | 14 | Frontal_Inf_Tri_R | AAL |
| 15 | -36 | 31 | -12 | Frontal_Inf_Orb_L | AAL |
| 16 | 41 | 32 | -12 | Frontal_Inf_Orb_R | AAL |
| 17 | -47 | -8 | 14 | Rolandic_Oper_L | AAL |
| 18 | 53 | -6 | 15 | Rolandic_Oper_R | AAL |
| 19 | -5 | 5 | 61 | Supp_Motor_Area_L | AAL |
| 20 | 9 | 0 | 62 | Supp_Motor_Area_R | AAL |
| 21 | -8 | 15 | -11 | Olfactory_L | AAL |
| 22 | 10 | 16 | -11 | Olfactory_R | AAL |
| 23 | -5 | 49 | 31 | Frontal_Sup_Medial_L | AAL |
| 24 | 9 | 51 | 30 | Frontal_Sup_Medial_R | AAL |
| 25 | -5 | 54 | -7 | Frontal_Med_Orb_L | AAL |
| 26 | 8 | 52 | -7 | Frontal_Med_Orb_R | AAL |
| 27 | -5 | 37 | -18 | Rectus_L | AAL |
| 28 | 8 | 36 | -18 | Rectus_R | AAL |
| 29 | -35 | 7 | 3 | Insula_L | AAL |
| 30 | 39 | 6 | 2 | Insula_R | AAL |
| 31 | -4 | 35 | 14 | Cingulum_Ant_L | AAL |
| 32 | 8 | 37 | 16 | Cingulum_Ant_R | AAL |
| 33 | -5 | -15 | 42 | Cingulum_Mid_L | AAL |
| 34 | 8 | -9 | 40 | Cingulum_Mid_R | AAL |
| 35 | -5 | -43 | 25 | Cingulum_Post_L | AAL |
| 36 | 7 | -42 | 22 | Cingulum_Post_R | AAL |
| 37 | 29 | -20 | -10 | Hippocampus_R | AAL |
| 38 | -25 | -21 | -10 | Hippocampus_L | AAL |
| 39 | -21 | -19 | -23 | ParaHippocampal_L | AAL |
| 40 | 21 | -19 | -23 | ParaHippocampal_R | AAL |
| 41 | -23 | -1 | -17 | Amygdala_L | AAL |
| 42 | 27 | 1 | -18 | Amygdala_R | AAL |
| 43 | -7 | -79 | 6 | Calcarine_L | AAL |
| 44 | 16 | -73 | 9 | Calcarine_R | AAL |
| 45 | -6 | -80 | 27 | Cuneus_L | AAL |

| | | | | | |
|---|---|---|---|---|---|
| 46 | 14 | -79 | 28 | Cuneus_R | AAL |
| 47 | -15 | -68 | -5 | Lingual_L | AAL |
| 48 | 16 | -67 | -4 | Lingual_R | AAL |
| 49 | -17 | -84 | 28 | Occipital_Sup_L | AAL |
| 50 | 24 | -81 | 31 | Occipital_Sup_R | AAL |
| 51 | -32 | -81 | 16 | Occipital_Mid_L | AAL |
| 52 | 37 | -80 | 19 | Occipital_Mid_R | AAL |
| 53 | -36 | -78 | -8 | Occipital_Inf_L | AAL |
| 54 | 38 | -82 | -8 | Occipital_Inf_R | AAL |
| 55 | -31 | -40 | -20 | Fusiform_L | AAL |
| 56 | 34 | -39 | -20 | Fusiform_R | AAL |
| 57 | -42 | -23 | 49 | Postcentral_L | AAL |
| 58 | 41 | -25 | 53 | Postcentral_R | AAL |
| 59 | -23 | -60 | 59 | Parietal_Sup_L | AAL |
| 60 | 26 | -59 | 62 | Parietal_Sup_R | AAL |
| 61 | -43 | -46 | 47 | Parietal_Inf_L | AAL |
| 62 | 46 | -46 | 50 | Parietal_Inf_R | AAL |
| 63 | -56 | -34 | 30 | SupraMarginal_L | AAL |
| 64 | 58 | -32 | 34 | SupraMarginal_R | AAL |
| 65 | -44 | -61 | 36 | Angular_L | AAL |
| 66 | 46 | -60 | 39 | Angular_R | AAL |
| 67 | -7 | -56 | 48 | Precuneus_L | AAL |
| 68 | 10 | -56 | 44 | Precuneus_R | AAL |
| 69 | -8 | -25 | 70 | Paracentral_Lobule_L | AAL |
| 70 | 7 | -32 | 68 | Paracentral_Lobule_R | AAL |
| 71 | -11 | 11 | 9 | Caudate_L | AAL |
| 72 | 15 | 12 | 9 | Caudate_R | AAL |
| 73 | -24 | 4 | 2 | Putamen_L | AAL |
| 74 | 28 | 5 | 2 | Putamen_R | AAL |
| 75 | -18 | 0 | 0 | Pallidum_L | AAL |
| 76 | 21 | 0 | 0 | Pallidum_R | AAL |
| 77 | -11 | -18 | 8 | Thalamus_L | AAL |
| 78 | 13 | -18 | 8 | Thalamus_R | AAL |
| 79 | -42 | -19 | 10 | Heschl_L | AAL |
| 80 | 46 | -17 | 10 | Heschl_R | AAL |
| 81 | -53 | -21 | 7 | Temporal_Sup_L | AAL |
| 82 | 58 | -22 | 7 | Temporal_Sup_R | AAL |
| 83 | -40 | 15 | -20 | Temporal_Pole_Sup_L | AAL |
| 84 | 48 | 15 | -17 | Temporal_Pole_Sup_R | AAL |
| 85 | -56 | -34 | -2 | Temporal_Mid_L | AAL |
| 86 | 57 | -37 | -1 | Temporal_Mid_R | AAL |
| 87 | -36 | 15 | -34 | Temporal_Pole_Mid_L | AAL |
| 88 | 44 | 15 | -32 | Temporal_Pole_Mid_R | AAL |
| 89 | -50 | -28 | -23 | Temporal_Inf_L | AAL |
| 90 | 54 | -31 | -22 | Temporal_Inf_R | AAL |
| 91 | -35 | -67 | -29 | Cerebellum_Crus1_L | AAL |
| 92 | 38 | -67 | -30 | Cerebellum_Crus1_R | AAL |
| 93 | -28 | -73 | -38 | Cerebellum_Crus2_L | AAL |

| | | | | | |
|---|---|---|---|---|---|
| 94 | 33 | -69 | -40 | Cerebellum_Crus2_R | AAL |
| 95 | -8 | -37 | -19 | Cerebellum_3_L | AAL |
| 96 | 13 | -34 | -19 | Cerebellum_3_R | AAL |
| 97 | -14 | -43 | -17 | Cerebellum_4_5_L | AAL |
| 98 | 18 | -43 | -18 | Cerebellum_4_5_R | AAL |
| 99 | -22 | -59 | -22 | Cerebellum_6_L | AAL |
| 100 | 26 | -58 | -24 | Cerebellum_6_R | AAL |
| 101 | -31 | -60 | -45 | Cerebellum_7b_L | AAL |
| 102 | 34 | -63 | -48 | Cerebellum_7b_R | AAL |
| 103 | -25 | -55 | -48 | Cerebellum_8_L | AAL |
| 104 | 26 | -56 | -49 | Cerebellum_8_R | AAL |
| 105 | -10 | -49 | -46 | Cerebellum_9_L | AAL |
| 106 | 10 | -49 | -46 | Cerebellum_9_R | AAL |
| 107 | -22 | -34 | -42 | Cerebellum_10_L | AAL |
| 108 | 27 | -34 | -41 | Cerebellum_10_R | AAL |
| 109 | 2 | -39 | -20 | Vermis_1_2 | AAL |
| 110 | 2 | -40 | -11 | Vermis_3 | AAL |
| 111 | 2 | -52 | -6 | Vermis_4_5 | AAL |
| 112 | 2 | -67 | -15 | Vermis_6 | AAL |
| 113 | 2 | -72 | -25 | Vermis_7 | AAL |
| 114 | 2 | -64 | -34 | Vermis_8 | AAL |
| 115 | 2 | -55 | -35 | Vermis_9 | AAL |
| 116 | 1 | -46 | -32 | Vermis_10 | AAL |
| 117 | -34 | -76 | 49 | Parietal_Inf_Lobe_L | [4] |
| 118 | -4 | 28 | 42 | Superior_Medial_Gyrus_L | [5] |
| 119 | 28 | -10 | -37 | Enthorinal_cortex_R | [6] |
| 120 | -23 | -8 | -38 | Enthorinal_cortex_L | [6] |
| 121 | -1 | -17 | 55 | Posterior_Medial_Frontal_L (SMA_L) | [7] |
| 122 | 8 | -17 | 57 | Posterior_Medial_Frontal_R (SMA_R) | [7] |
| 123 | -17 | -58 | -38 | Dentate_Nucleus_L | [8] |
| 124 | 14 | -61 | -38 | Dentate_Nucleus_R | [8] |
| 125 | 11 | -12 | 0 | Subthalamic_Nucleus_R | [9] |
| 126 | -8 | -13 | -7 | Subthalamic_Nucleus_L | [9] |

**Table 5.** Full connectome node list (116 AAL nodes (not highlighted); 10 additional nodes inspired by literature (in grey). The MNI coordinates of additional nodes derived/were checked with the Anatomy toolbox (SPM).

**Table 6. The node list of the wideband rsFC network for the induction effect (Rest 3 vs. Rest 2) in the boost time window**

| Number | Node's weight | Region |
|---|---|---|
| 1 | 9.0 | Precentral_L |
| 2 | 7.2 | Frontal_Inf_Oper_R |
| 3 | 3.9 | Rolandic_Oper_R |

| | | |
|---|---|---|
| 4 | 19.4 | Supp_Motor_Area_R |
| 5 | 25.9 | Insula_L |
| 6 | 7.3 | Cingulum_Mid_R |
| 7 | 3.9 | Cingulum_Post_R |
| 8 | 3.8 | Occipital_Sup_L |
| 9 | 11.6 | Occipital_Mid_L |
| 10 | 7.4 | Occipital_Mid_R |
| 11 | 8.6 | Postcentral_L |
| 12 | 7.8 | Parietal_Inf_L |
| 13 | 3.9 | Angular_R |
| 14 | 7.4 | Precuneus_L |
| 15 | 3.8 | Precuneus_R |
| 16 | 3.6 | Paracentral_Lobule_L |
| 17 | 7.4 | Paracentral_Lobule_R |
| 18 | 3.6 | Putamen_R |
| 19 | 7.2 | Pallidum_R |
| 20 | 3.6 | Heschl_L |
| 21 | 31.3 | Temporal_Sup_L |
| 22 | 39.8 | Temporal_Mid_L |
| 23 | 10.8 | Cerebellum_Crus2_L |
| 24 | 10.8 | Cerebellum_6_L |
| 25 | 7.2 | Cerebellum_6_R |
| 26 | 7.2 | Vermis_7 |
| 27 | 7.7 | Posterior_Medial_Frontal _R (SMA_R) |
| 28 | 26.8 | Dentate_Nucleus_L |

**Table 6.** The node list of the wideband resting-state functional connectivity (rsFC) analysis for the induction effect (Rest 3 vs. Rest 2) in the boost time window.

### 4. Resting-state functional connectivity 5-band data

To reveal band-specific rsFC changes induced by motor learning, we computed the adjacency matrix (126 by 126) for five distinct frequency bands: delta (δ 1–4 Hz), theta (θ band: 4–8 Hz), alpha (α band: 8–13 Hz), beta (β band: 13–30 Hz) and gamma (γ band: 30–45 Hz). Next, we performed a statistical within-group comparison of inter-regional connectivity differences as described for the wideband in the previous section. The results of this analysis are illustrated in Figures 4 - 10.

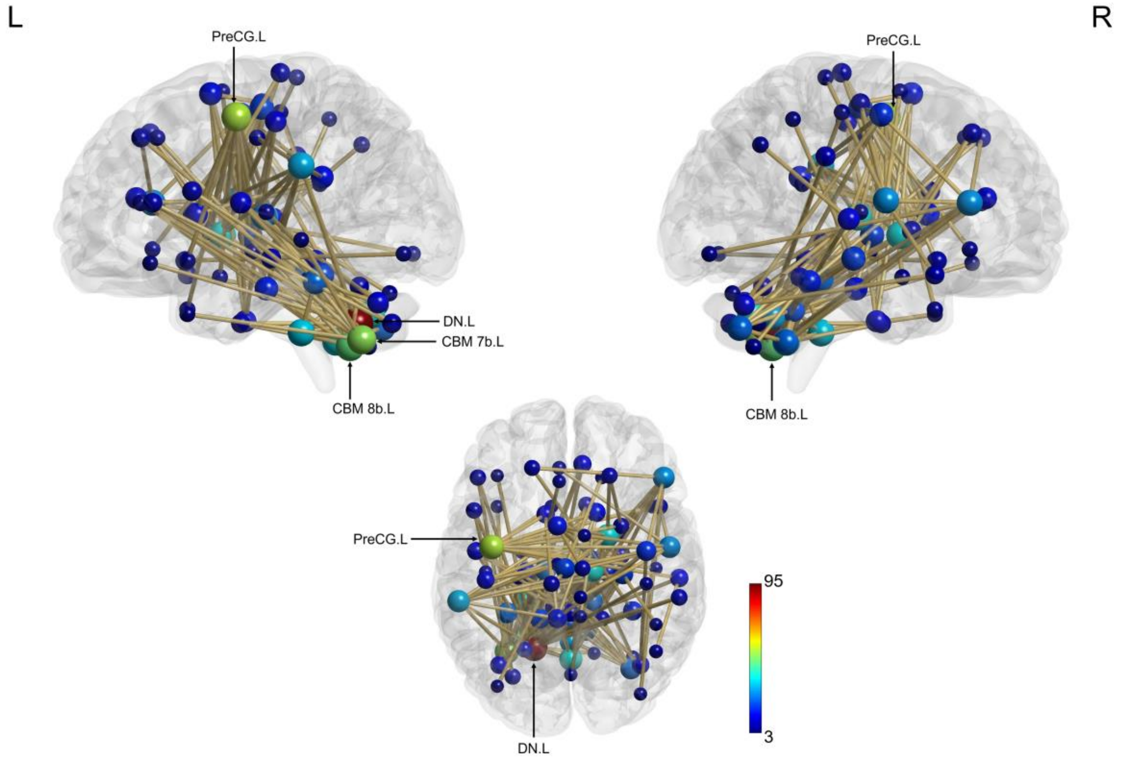


**Figure 4.** Neural network representing increased connectivity in post-learning session (Rest 2 vs. Rest 1) in the theta band. Nodes are scaled according to their weight (the sum of all edges connected to the node). Significant edges are represented as interconnecting lines between 126 connectome seed regions. Abbreviations: PreCG – Precentral gyrus; DN – Dentate nucleus. L – left; R – right.

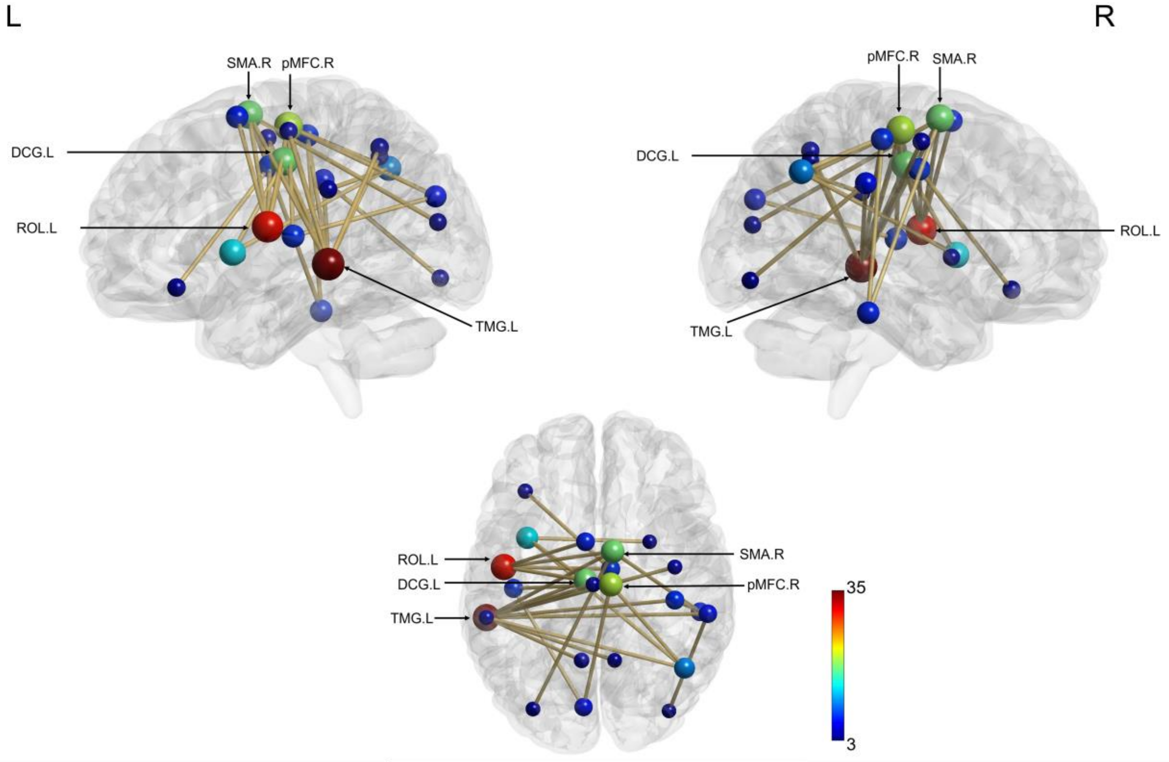


**Figure 5.** Neural network representing the increased connectivity in the alpha band as a result of induction effect (re-introduction to the task) during the boost period. Nodes are scaled according to their weight (the

sum of all edges connected to the node). Significant edges are represented as interconnecting lines between 126 connectome seed regions. Abbreviations: ROL – Rolandic operculum; TMG – Temporal middle gyrus; pMFC – posterior Medial frontal cortex; SMA - Supplementary motor area; DCG – Median cingulate and paracingulate gyri; L – left; R – right.

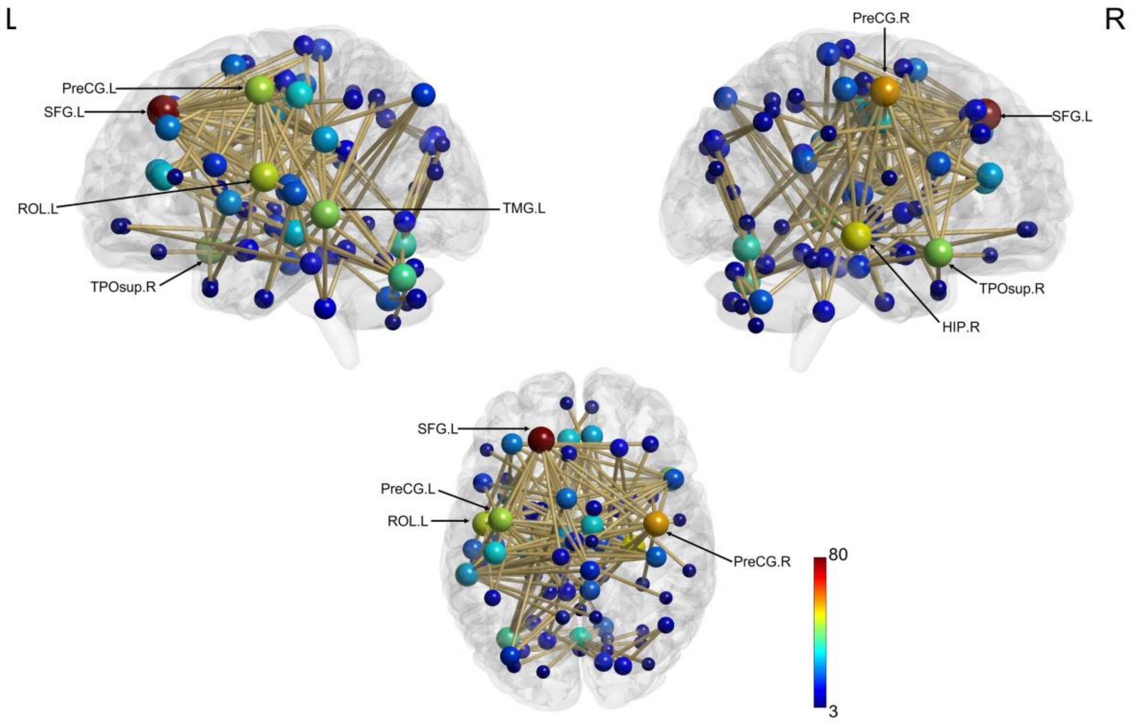


**Figure 6.** Neural network representing the increased connectivity in the beta band as a result of induction effect (re-introduction to the task) during the boost period. Nodes are scaled according to their weight (the sum of all edges connected to the node). Significant edges are represented as interconnecting lines between 126 connectome seed regions. Abbreviations: SFG - Superior frontal gurus; PreCG – Precentral gyrus; ROL – Rolandic operculum; L – left; R – right.

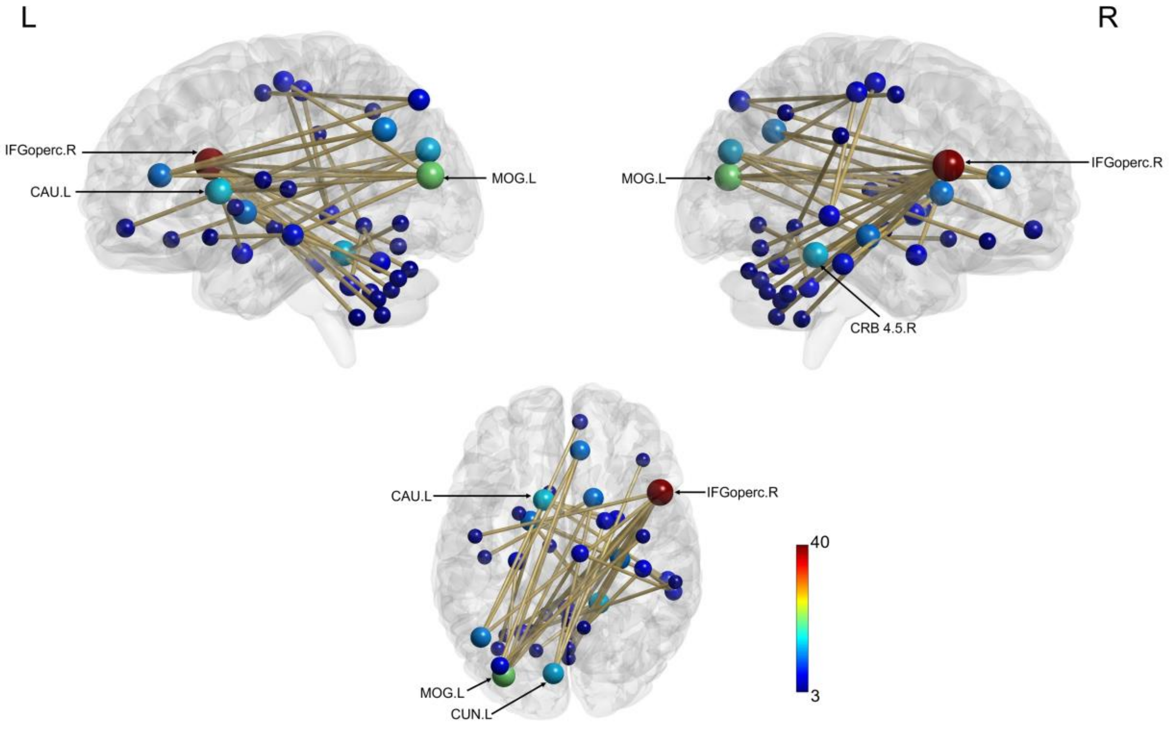


**Figure 7.** Neural network representing the increased connectivity in the delta band as a result of induction effect (re-introduction to the task) during the boost period. Nodes are scaled according to their weight (the sum of all edges connected to the node). Significant edges are represented as interconnecting lines between 126 connectome seed regions. Abbreviations: IFGoperc – Inferior frontal gyrus, opercular part; MOG – Middle occipital gyrus; CAU - Caudate nucleus; CUN – Cuneus; L – left; R – right.

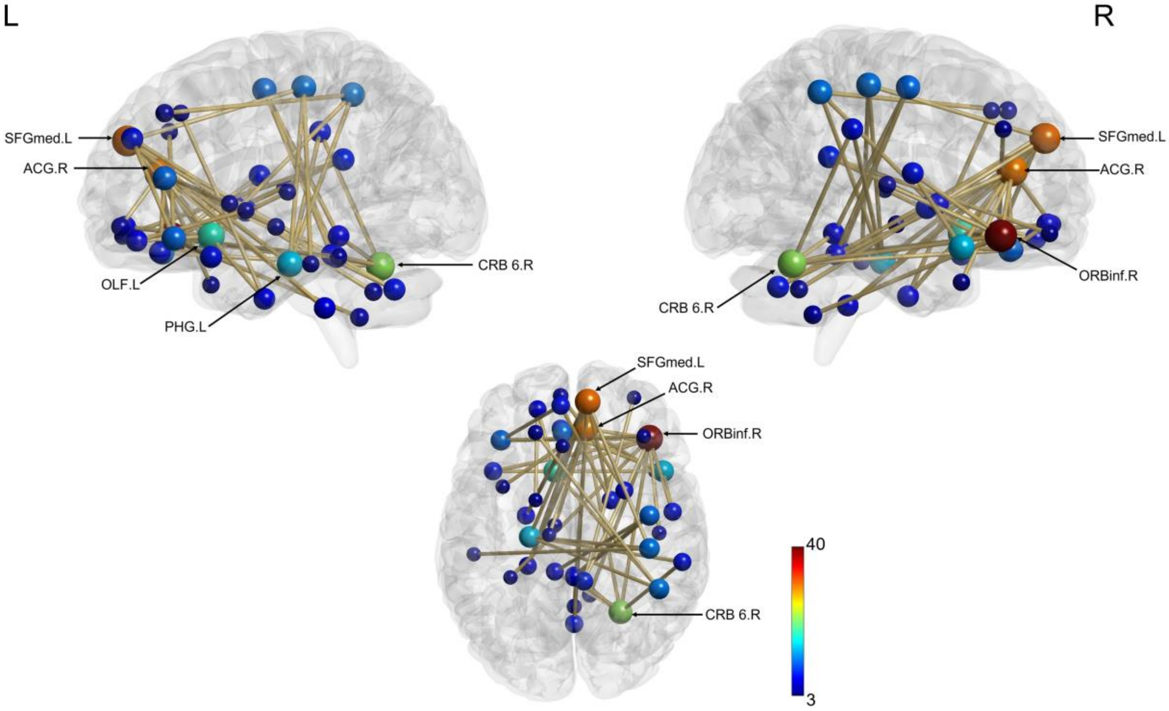


**Figure 8.** Neural network representing the increased connectivity in gamma band as a result of induction effect (re-introduction to the task) during the boost period. Nodes are scaled according to their weight (the sum of all edges connected to the node). Significant edges are represented as interconnecting lines between 126 connectome seed regions. Abbreviations: ORBinf – Inferior frontal gyrus, orbital part; ACG –

Anterior cingulate and paracingulate gyri; SFGmed – Superior frontal gyrus, medial; CRB 6 – Cerebellum lobe 6; L – left; R – right.

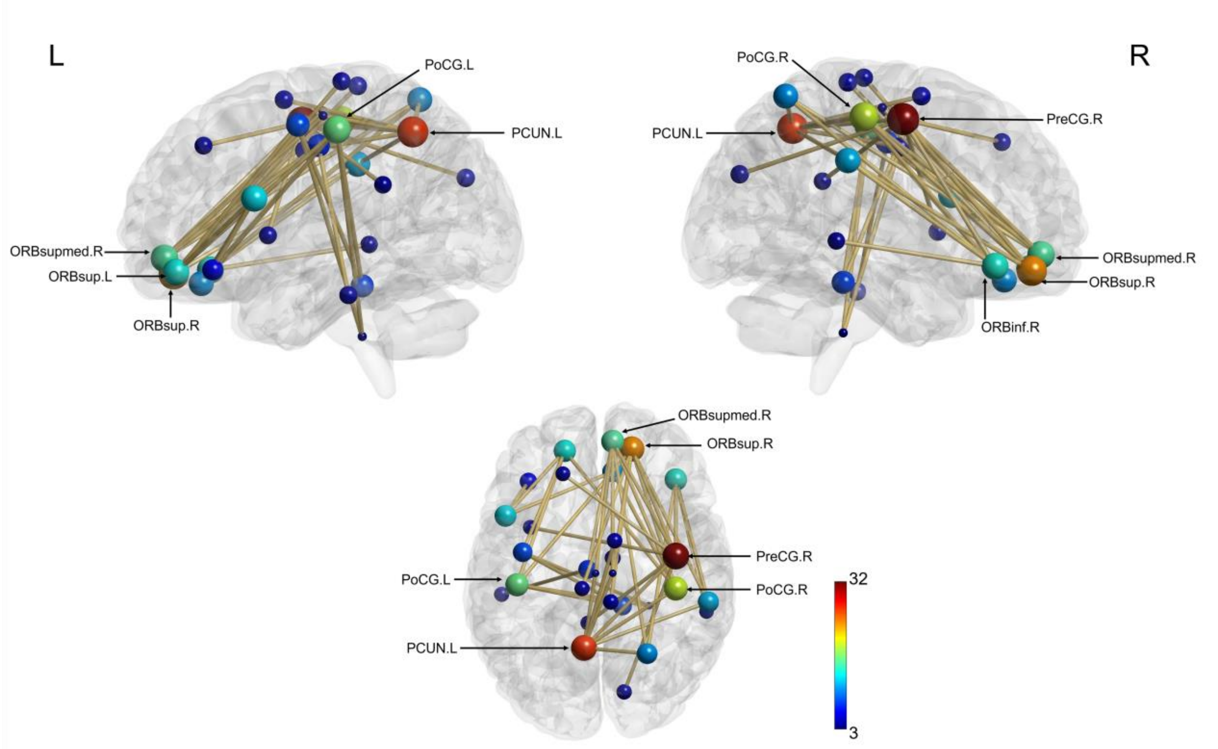


**Figure 9.** Neural network representing the increased connectivity in alpha band as a result of induction effect (re-introduction to the task) during the next day period. Nodes are scaled according to their weight (the sum of all edges connected to the node). Significant edges are represented as interconnecting lines between 126 connectome seed regions. Abbreviations: PreCG – Precental gyrus; PCUN - Precuneus; ORBsup – Superior frontal gyrus, orbital part; PoCG – Postcentral gyrus; ORBsupmed – Superior frontal gyrus, medial orbital; L – left; R – right.

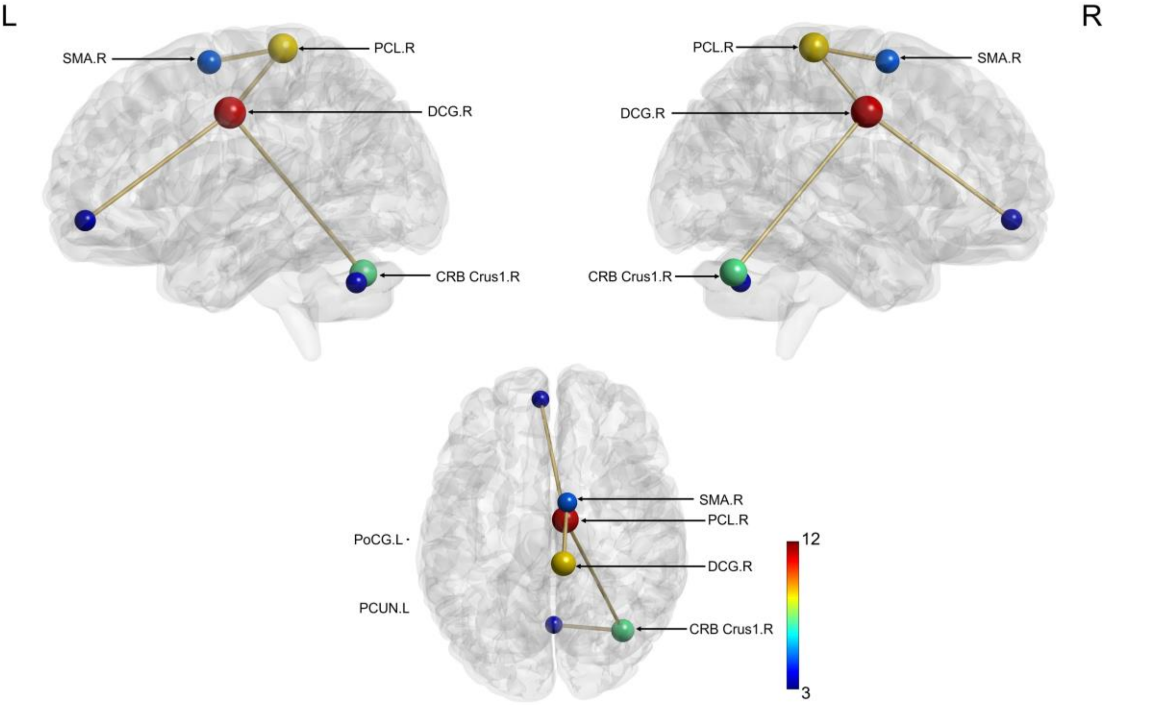

**Figure 10.** Neural network representing the increased connectivity in gamma band as a result of induction effect (re-introduction to the task) during the next day period. Nodes are scaled according to their weight (the sum of all edges connected to the node). Significant edges are represented as interconnecting lines between 126 connectome seed regions. Abbreviations: PCL – Paracentral lobule; DCG – Median cingulate and paracingulate gyri; CRB Crus 1 – Cerebellum Crus 1; SMA – Supplementary motor area; L – left; R – right.

### 5. Correlational analysis between connectivity and behavioural indices

Correlational analyses between each resting-state RS session and Best Motor Performance Index (BMP) index highlighted a positive correlation between BMP and RS 3 connectivity (boost period, induced). Table 7 shows the list of nodes for the revealed resting-state functional connectivity network.

**Table 7. Nodes list of the neural network which positively correlated with BMP**

| Number | Node's weight | Region |
|---|---|---|
| 1 | 19.8 | Frontal_Sup_R |
| 2 | 11.3 | Frontal_Mid_Orb_L |
| 3 | 28.7 | Frontal_Mid_Orb_R |
| 4 | 9.1 | Rolandic_Oper_R |
| 5 | 11.6 | Frontal_Med_Orb_L |
| 6 | 42.3 | Frontal_Med_Orb_R |
| 7 | 3.9 | Cingulum_Ant_L |
| 8 | 3.8 | ParaHippocampal_R |
| 9 | 3.6 | Amygdala_R |
| 10 | 3.9 | Fusiform_R |
| 11 | 3.6 | Parietal_Sup_R |
| 12 | 3.9 | Parietal_Inf_R |
| 13 | 11.0 | Angular_L |
| 14 | 12.1 | Angular_R |
| 15 | 3.5 | Paracentral_Lobule_L |
| 16 | 4.3 | Caudate_L |
| 17 | 7.4 | Thalamus_L |
| 18 | 8.1 | Heschl_R |
| 19 | 12.3 | Temporal_Sup_R |
| 20 | 7.9 | Temporal_Inf_R |
| 21 | 3.8 | Cerebellum_3_R |
| 22 | 3.9 | Cerebellum_4_5_R |
| 23 | 4.0 | Enthorinal_cortex_R |
| 24 | 3.5 | Subthalamic_Nucleus_R |

**Table 7.** Nodes list of the neural network which positively correlated with Best Motor Performance (BMP).

Correlational analysis between rsFC and the Learning index (LI) revealed significant positive correlations only for RS 5 (induced RS, silent period). Table 8 lists the nodes from the emerged rsFC network.

**Table 8. Nodes list of the neural network which positively correlated with the Learning index**

| Number | Node's weight | Region |
|---|---|---|
| 1 | 9.2 | Frontal_Mid_R |
| 2 | 3.8 | Frontal_Inf_Oper_R |
| 3 | 3.8 | Frontal_Inf_Tri_R |
| 4 | 3.6 | Frontal_Inf_Orb_L |
| 5 | 10.7 | Olfactory_R |
| 6 | 14.6 | Insula_L |
| 7 | 14.6 | Insula_R |
| 8 | 11.5 | Hippocampus_R |
| 9 | 11.0 | Amygdala_R |
| 10 | 3.5 | Calcarine_L |
| 11 | 3.8 | Calcarine_R |
| 12 | 3.6 | Cuneus_L |
| 13 | 7.1 | Cuneus_R |
| 14 | 7.1 | Lingual_R |
| 15 | 3.7 | Occipital_Inf_R |
| 16 | 14.7 | Fusiform_R |
| 17 | 14.8 | Caudate_L |
| 18 | 3.5 | Caudate_R |
| 19 | 11.2 | Putamen_R |
| 20 | 19.9 | Thalamus_R |
| 21 | 8.5 | Temporal_Pole_Sup_L |
| 22 | 18.9 | Temporal_Pole_Sup_R |
| 23 | 12.0 | Temporal_Pole_Mid_L |
| 24 | 14.4 | Cerebellum_Crus1_L |
| 25 | 7.3 | Cerebellum_Crus1_R |
| 26 | 7.3 | Cerebellum_Crus2_R |
| 27 | 3.5 | Cerebellum_6_L |
| 28 | 3.8 | Cerebellum_8_L |
| 29 | 8.2 | Cerebellum_9_L |
| 30 | 8.2 | Cerebellum_9_R |
| 31 | 3.7 | Vermis_3 |
| 32 | 11.7 | Vermis_4_5 |
| 33 | 3.6 | Vermis_6 |
| 34 | 7.7 | Vermis_10 |
| 35 | 3.8 | Superior_Medial_Gyrus_L |
| 36 | 3.7 | Dentate_Nucleus_L |

**Table 8.** Nodes list of the neural network which positively correlated with the Learning index (LI).

### 6. State networks connectivity strength

Using HMM power maps, we extracted three sets of nodes (state networks) representing HMM States 3, 7, and 8 (see main text results). To quantify the connectivity strength of these state networks, we calculated the sum of all power envelope correlation values for each node within a given network. Next, we divided this sum by the number of nodes within the network to obtain a normalised measure of connectivity strength. This procedure enabled us to generate a single value for each subject in each resting-state session, which was used in a subsequent correlational analysis with behavioural measures using JASP version 0.16.2, JASP Team (2022). The results disclosed positive correlations between BMP and State 7 (Frontal/Cuneus) network. Table 9 lists the nodes comprising the network.

**Table 9. The node list of the HMM state 7 (Frontal/Cuneus) network**

| Number | Region |
|---|---|
| 1 | Frontal_Sup_L |
| 2 | Frontal_Sup_R |
| 3 | Frontal_Sup_Orb_L |
| 4 | Frontal_Sup_Orb_R |
| 5 | Frontal_Mid_L |
| 6 | Frontal_Mid_R |
| 7 | Frontal_Mid_Orb_L |
| 8 | Frontal_Mid_Orb_R |
| 9 | Frontal_Inf_Tri_L |
| 10 | Frontal_Inf_Tri_R |
| 11 | Frontal_Inf_Orb_R |
| 12 | Frontal_Sup_Medial_L |
| 13 | Frontal_Sup_Medial_R |
| 14 | Frontal_Med_Orb_L |
| 15 | Frontal_Med_Orb_R |
| 16 | Rectus_L |
| 17 | Rectus_R |
| 18 | Cingulum_Ant_R |

**Table 9.** The node list of the HMM state 7 (Frontal/Cuneus) network.

### 7. Fast and slow neural dynamics relationships

To reveal the relationship between fast and slow neural dynamics, we performed a correlational analysis between HMM temporal parameters and state networks' rsFC using the NBS toolbox. Table 10 illustrates the correlational outcomes for State 8.

**Table 10. Correlations between HMM temporal parameters (TP) and HMM State 8 (Cuneus/Sensorimotor) network**

| Rest / TP | FO | MLT | NO | MIL |
|---|---|---|---|---|
| Rest 1 | ***p* = .023#**<br>(1 edge, 2 nodes) | *p* = 1.00 (n.s) | *p* = 1.00 (n.s) | ***p* = .022#**<br>(1 edge, 2 nodes) |
| Rest 2 | *p* = 1.00 (n.s) | *p* = 1.00 (n.s) | *p* = 1.00 (n.s) | ***p* = .03#**<br>(1 edge, 2 nodes) |
| Rest 3 | *p* = 1.00 (n.s) | *p* = 1.00 (n.s) | ***p* = .01#**<br>(2 edges, 3 nodes) | *p* = 1.00 (n.s) |
| Rest 4 | *p* = 1.00 (n.s) | *p* = 1.00 (n.s) | *p* = 1.00 (n.s) | *p* = 1.00 (n.s) |
| Rest 5 | *p* = 1.00 (n.s) | *p* = 1.00 (n.s) | *p* = 1.00 (n.s) | *p* = 1.00 (n.s) |
| Rest 6 | *p* = 1.00 (n.s) | *p* = 1.00 (n.s) | *p* = 1.00 (n.s) | *p* = 1.00 (n.s) |
| Rest 7 | *p* = 1.00 (n.s) | *p* = 1.00 (n.s) | *p* = 1.00 (n.s) | *p* = 1.00 (n.s) |

**Table 10.** Correlations between HMM temporal parameters (TP) and HMM State 8 (Cuneus/Sensorimotor) network. Spearman's rho and *p*-values (#*p* ≤ .05, uncorrected).

[9] B. U. Forstmann *et al.*, "Cortico-subthalamic white matter tract strength predicts interindividual

efficacy in stopping a motor response," *Neuroimage*, vol. 60, no. 1, pp. 370–375, 2012, doi: 10.1016/j.neuroimage.2011.12.044.
